# Genomic correlates of hyperthermostability revisited: a large-scale validation across 1,963 microbial genomes

**DOI:** 10.64898/2026.04.18.719035

**Authors:** Karsten Suhre

## Abstract

In 2003, Suhre and Claverie demonstrated that the difference between the fraction of charged amino acids and the fraction of polar uncharged amino acids in a proteome (the CvP-bias) was the single genomic feature that most strongly discriminates hyperthermophilic microorganisms from their mesophilic and thermophilic counterparts. The original analysis was based on 71 completely sequenced genomes available at the time. Here, using modern genome databases — specifically the Bacterial and Viral Bioinformatics Resource Center (BV-BRC) for curated optimal growth temperature (OGT) metadata and the NCBI RefSeq FTP archive for sequence data — the same analysis is repeated at approximately 28-fold larger scale, covering 1,963 bacterial and archaeal genomes (103 hyperthermophiles, 409 thermophiles, 1,451 mesophiles). The original finding is confirmed with high statistical confidence: CvP-bias is elevated in hyperthermophiles (mean 11.1 ± 3.1%) relative to mesophiles (5.0 ± 2.0%; ANOVA F = 496, *p* < 10^−170^), with a large effect size (Cohen’s *d* = 2.32) and area under the receiver operating characteristic curve of 0.94 for binary hyper/meso classification. Principal component analysis confirms that the first principal component, explaining 47% of variance, is loaded by CvP-bias as a major contributor, separating hyperthermophiles from other organisms. These results establish that the CvP-bias signal identified in 2003 is not an artifact of small sample size but a genuine, robust property of hyperthermophilic proteomes.

## 1. Introduction

The adaptation of microorganisms to extreme temperatures requires molecular stabilization of proteins, nucleic acids, and membranes under conditions that would denature or inactivate their mesophilic homologues. Hyperthermophiles — organisms with an optimal growth temperature (OGT) above 80°C — include representatives of both the Archaea (e.g., *Pyrococcus, Methanocaldococcus, Archaeoglobus, Sulfolobus*) and Bacteria (e.g., *Aquifex, Thermotoga*), and are regarded as among the most ancient lineages of life [1,2]. Understanding the molecular basis of thermostability has implications for protein engineering, industrial enzymology, and the search for life in extreme environments.

At the proteome level, multiple compositional biases have been proposed as correlates of thermostability. Elevated fractions of charged residues (particularly Asp, Glu, Lys, Arg, His) relative to polar uncharged residues (Asn, Gln, Ser, Thr) — the CvP-bias — were identified by several groups as a characteristic feature of hyperthermophile proteins [3,4]. Additionally, the average isoelectric point (pI) of the proteome and a dinucleotide deviation index S* computed from coding sequences have been proposed as complementary discriminators [5,6].

In 2003, Suhre and Claverie conducted a systematic analysis of all 71 completely sequenced microbial genomes then available, quantifying CvP-bias, mean proteome pI, and dinucleotide S* for each organism [5]. A principal component analysis (PCA) of these three features revealed that the first principal component, which captured the majority of between-organism variance, was dominated by the CvP-bias loading, placing all nine hyperthermophilic genomes at one extreme. The finding was summarized as: CvP-bias is the sole genomic correlate of hyperthermostability that can be detected from complete genome sequences.

In the years following the 2003 study, the genomic correlates of thermal adaptation attracted considerable further investigation from several directions. Zeldovich, Berezovsky, and Shakhnovich [13] analysed 204 complete proteomes spanning the full biological temperature range from −10°C to 110°C and identified a seven-amino-acid composition index — the IVYWREL set (Ile, Val, Tyr, Trp, Arg, Glu, Leu) — whose proteome-wide frequency correlates with OGT at *r* = 0.93, yielding a near-linear predictor (Topt ≈ 937FIVYWREL − 335). Although IVYWREL is physically heterogeneous, mixing aliphatic, aromatic, and charged residues, it became a widely adopted benchmark for genome-based temperature inference. At the nucleotide level, GC content of rRNA stem-loop structures, codon composition, and genomic GC content have all been reported as correlates of OGT [14,15], while bacterial genome size decreases with growth temperature in a pattern interpreted as selective streamlining under thermal pressure [16]. Asymmetry between hot and cold adaptation has also been noted: the amino acid compositional shifts in psychrophiles are not simply the mirror image of those in hyperthermophiles [17], suggesting that different selective forces operate at the two temperature extremes. Machine learning approaches incorporating hundreds of genome-derived features — codon frequencies, amino acid composition vectors, dinucleotide spectra — have since achieved continuous OGT regression with root-mean-square errors below 5°C [18,19], substantially outperforming single-index metrics quantitatively, though at the cost of mechanistic interpretability. Most recently, a large-scale evolutionary analysis of nearly 3,000 prokaryotes found that OGT predictability from genome composition is itself modulated by the rate of evolutionary temperature shifts, with lineages that recently colonised cold environments being systematically less predictable from sequence composition alone [20].

The intervening two decades have thus seen an enormous expansion both in the number of sequenced genomes and in the sophistication of OGT predictors. This provides an opportunity to re-examine whether the CvP-bias signal of the 2003 study reflects a genuine biological phenomenon or was inflated by the small and phylogenetically restricted genome set then available.

The present study repeats the 2003 analysis using modern data, with no modification to the original computational methodology, at approximately 28-fold larger scale. The analysis was performed entirely using open-source tools and scripted in Python, with workflow automation provided by Claude Code (Anthropic), a large language model-based programming assistant.

## 2. Materials and Methods

### 2.1 Genome selection and metadata

Microbial genomes with curated temperature classification were retrieved from the Bacterial and Viral Bioinformatics Resource Center (BV-BRC; bv-brc.org) [7] using the BV-BRC REST API with RQL (Resource Query Language) queries against the `/api/genome/` endpoint. The fields retrieved included genome identifier, genome name, taxonomic information, assembly accession (GCF/GCA), OGT, and temperature range classification. Queries were made separately for each temperature category: Mesophilic, Thermophilic, and Hyperthermophilic. Psychrophilic organisms were excluded to focus on the heat-adapted spectrum, consistent with the original analysis.

Organisms were classified as follows: hyperthermophile (OGT > 80°C or BV-BRC classification “Hyperthermophilic”), thermophile (50–80°C or “Thermophilic”), and mesophile (< 50°C or “Mesophilic”). Only Bacteria and Archaea were retained; Eukaryotes were excluded.

To obtain genome sequence data, BV-BRC assembly accession numbers were mapped to NCBI RefSeq FTP paths using the bulk assembly summary files (`assembly_summary.txt`) for bacteria and archaea downloaded from the NCBI FTP archive (ftp.ncbi.nlm.nih.gov/genomes/refseq/). This approach provided FTP paths for 978,000 assemblies without the need for individual Entrez API queries. BV-BRC accessions were matched against NCBI entries with and without version suffix, and GCF/GCA prefix interconversion was attempted where direct matches failed.

From the matched set (7,394 of 8,730 BV-BRC annotated genomes), the final dataset was constructed by including all hyperthermophilic (n = 104) and all thermophilic (n = 413) genomes with FTP paths, supplemented by a stratified random sample of up to 1,500 mesophilic genomes sampled proportionally by phylum to reduce taxonomic bias. This yielded 2,017 candidate genomes. Protein FASTA files (`*_protein.faa.gz`) and coding sequence FASTA files (`*_cds_from_genomic.fna.gz`) were downloaded in parallel using 10 concurrent threads. Files were validated as non-empty, correctly gzip-compressed FASTA format before acceptance. Of the 2,017 candidates, 1,965 were successfully downloaded (97.4%); after metric computation, 1,963 genomes yielded valid results and were included in the final analysis.

### 2.2 CvP-bias

CvP-bias was computed as described by Suhre and Claverie [5] and, independently, by other groups [3,4]. All protein sequences from the protein FASTA file were concatenated, and the frequencies of each amino acid were calculated over the concatenated sequence, excluding ambiguous characters (B, J, O, U, X, Z) and gap characters. CvP-bias is then:

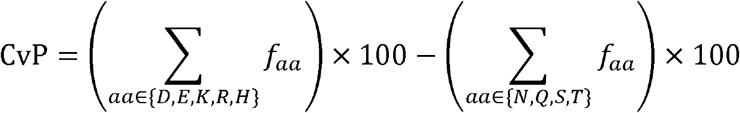

where *f*aa is the frequency of amino acid aa in the concatenated proteome.

### 2.3 Mean proteome isoelectric point

The isoelectric point (pI) of each individual protein was calculated using the `ProteinAnalysis.isoelectric_point()` method from BioPython v1.87 [8], which implements the standard Henderson–Hasselbalch iterative method with the pK values of Lehninger. Proteins shorter than three residues after removal of ambiguous characters were excluded. The mean proteome pI was defined as the arithmetic mean of the per-protein pI values.

### 2.4 Dinucleotide statistical index S*

The dinucleotide deviation index S* was computed as described by Karlin and Mrazek [6] and applied to thermophile analysis by Suhre and Claverie [5]. All coding sequences were pooled, and mono- and dinucleotide frequencies were counted over the complete nucleotide content of the coding FASTA. The relative dinucleotide abundance was defined as:

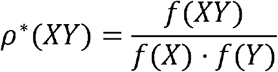

where *f*(XY) is the observed frequency of the dinucleotide XY and *f*(X), *f*(Y) are the marginal mononucleotide frequencies. The index S* was then:

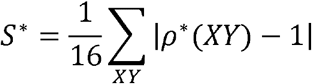

For organisms lacking a valid CDS FASTA file (52 genomes), S* was recorded as missing.

### 2.5 Statistical analysis and principal component analysis

Between-group differences in each metric were assessed by one-way ANOVA across all three classes and by two-sample *t*-tests for pairwise comparisons (Welch’s variant, unequal variance). Effect size was quantified by Cohen’s *d*. Binary classification performance of CvP-bias alone (hyperthermophile vs. mesophile) was evaluated using the area under the receiver operating characteristic curve (AUC-ROC).

PCA was performed on the three-metric matrix (CvP, mean pI, S*) after per-feature standardization (zero mean, unit variance) using `sklearn.decomposition.PCA` from scikit-learn [9]. Only genomes with complete data for all three metrics were included in the PCA (n = 1,911). Loadings (principal component coefficients) were extracted to identify the features contributing most to each principal component. A biplot was constructed by overlaying scaled loading vectors on the scatter plot of PC1 vs. PC2 scores, color-coded by temperature class.

### 2.6 Reproducibility

All scripts, data download procedures, and analyses were implemented in Python 3 and are provided as Supplementary Data. The workflow was designed and coded interactively using Claude Code (Anthropic, claude-sonnet-4-6 model). Raw genome sequence files are not redistributed but are freely downloadable from NCBI RefSeq FTP and BV-BRC; the download scripts and checkpoint files provided in Supplementary Data are sufficient to reconstruct the dataset.

## 3. Results

### 3.1 Dataset composition

The final dataset comprises 1,963 microbial genomes: 103 hyperthermophiles, 409 thermophiles, and 1,451 mesophiles (Table 1). The dataset is dominated by Bacteria (1,845 genomes), with Archaea representing 172 genomes. At the phylum level, the most abundant groups are Pseudomonadota (formerly Proteobacteria; 714 genomes), Bacillota (formerly Firmicutes; 489 genomes), and Actinomycetota (formerly Actinobacteria; 194 genomes). Hyperthermophilic organisms are predominantly represented by Thermoproteota (38 genomes, all Archaea) and Methanobacteriota (37 genomes), consistent with the known taxonomic distribution of hyperthermophiles. Median proteome sizes range from 2,005 proteins per genome in hyperthermophiles to 3,628 in mesophiles, reflecting the typically compact genomes of hyperthermophilic Archaea.

**Table 1.** Dataset composition.

| Class | Genomes (n) | Bacteria | Archaea | OGT available |
| --- | --- | --- | --- | --- |
| Hyperthermophile (> 80°C) | 103 | 29 | 74 | 89 |
| Thermophile (50–80°C) | 409 | 335 | 74 | 295 |
| Mesophile (< 50°C) | 1,451 | 1,481* | 24 | 676 |
| **Total** | **1,963** | **1,845** | **172** | **1,060** |

*Note: Kingdom assignments are from BV-BRC; not all genomes in the metrics file were matched to the metadata.*

Of the 1,060 genomes with numeric OGT records, the mean OGT was 82.9 ± 10.7°C for hyperthermophiles, 55.3 ± 12.3°C for thermophiles, and 32.6 ± 5.2°C for mesophiles, consistent with the expected temperature ranges for each class.

### 3.2 CvP-bias is strongly elevated in hyperthermophiles

Figure 1 shows the distribution of all three metrics (CvP-bias, mean pI, S) stratified by temperature class. The most striking pattern concerns CvP-bias: the distributions for hyperthermophiles, thermophiles, and mesophiles are well-separated, with progressive increase in CvP from mesophiles (4.98 ± 2.04%) through thermophiles (7.94 ± 3.25%) to hyperthermophiles (11.12 ± 3.14%; Table 2). One-way ANOVA yields F = 496.0 (p* < 5 × 10^−175^) across the three classes. All pairwise comparisons are highly significant: mesophiles vs. thermophiles (*t* = 22.4, *p* < 10^−98^), mesophiles vs. hyperthermophiles (*t* = 28.3, *p* < 10^−142^), and thermophiles vs. hyperthermophiles (*t* = 9.0, *p* < 10^−17^). The effect size for the hyperthermophile-vs.-mesophile contrast is large (Cohen’s *d* = 2.32). Seventy-eight percent of hyperthermophilic genomes have CvP-bias values exceeding the 95th percentile of the mesophilic distribution.

The organisms with the highest CvP-bias values in the entire dataset are archetypical hyperthermophiles: *Methanopyrus kandleri* AV19 (OGT 98°C; CvP = 17.72%), *Methanocaldococcus infernus* ME (OGT 85°C; CvP = 15.91%), and *Aquifex aeolicus* VF5 (OGT 96°C; CvP = 15.40%).

**Table 2.** Summary statistics for the three genomic metrics by temperature class.

| Metric | Hyperthermophile | Thermophile | Mesophile | ANOVA F | p |
| --- | --- | --- | --- | --- | --- |
| CvP-bias (%) | 11.12 ± 3.14 | 7.94 ± 3.25 | 4.98 ± 2.04 | 496.0 | < 10<sup>-170</sup> |
| Mean pI | 7.31 ± 0.31 | 7.17 ± 0.43 | 6.95 ± 0.50 | 54.6 | < 10<sup>-23</sup> |
| $S^*$ | 0.148 ± 0.035 | 0.163 ± 0.039 | 0.157 ± 0.037 | 8.1 | 3.1 × 10<sup>-4</sup> |

Values are mean ± SD. Sample sizes: hyperthermophile n = 103, thermophile n = 409, mesophile n = 1,451 (CvP, pI); $S^*$ calculated for genomes with CDS data (n = 1,911).

**Figure 1.**
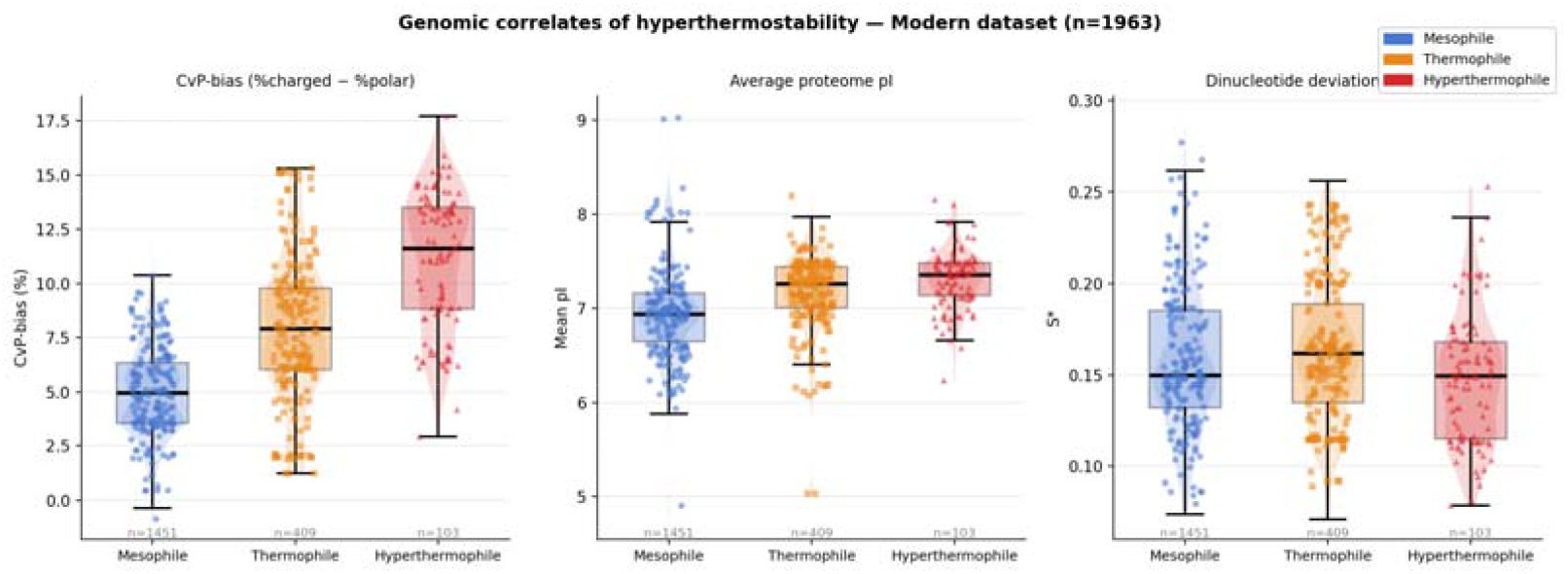
Distribution of CvP-bias (left), mean proteome pI (center), and dinucleotide index S* (right) for 1,963 microbial genomes stratified by temperature class (blue: mesophile, orange: thermophile, red: hyperthermophile). Box plots show median and interquartile range; violin plots indicate kernel density for n > 20; individual points (random subsample of up to 200 per class) are shown as jittered markers.

### 3.3 Mean proteome pI and dinucleotide S* show weaker associations

Mean proteome pI also differs significantly across classes (ANOVA F = 54.6, *p* < 10^−23^; Figure 1), with hyperthermophiles showing slightly higher mean pI (7.31 ± 0.31) than mesophiles (6.95 ± 0.50), and the thermophile group intermediate. However, the effect size is substantially smaller than for CvP-bias, and the distributions show considerable overlap.

The dinucleotide index S* shows a significant but modest overall ANOVA signal (F = 8.1, *p* = 3.1 × 10^−4^). Notably, hyperthermophiles have a *lower* mean S* (0.148 ± 0.035) than both thermophiles (0.163 ± 0.039) and mesophiles (0.157 ± 0.037). The pairwise test between hyperthermophiles and mesophiles is marginally significant in the negative direction (*t* = −2.4, *p* = 0.016). This pattern is discussed further in Section 4.

### 3.4 CvP-bias alone classifies hyperthermophiles with high accuracy

Using CvP-bias as the sole predictor in a binary classification of hyperthermophile vs. mesophile, the area under the receiver operating characteristic curve (AUC-ROC) is 0.941. This confirms that a single proteome-wide compositional metric, requiring no structural information, no gene annotation, and no comparative genomics, achieves near-optimal separation between temperature classes at a scale 28-fold larger than the original study.

### 3.5 CvP-bias correlates with optimal growth temperature

Among the 1,060 genomes with numeric OGT data, the Pearson correlation between CvP-bias and OGT is *r* = 0.691 (*p* < 10^−150^; n = 1,060). The scatter plot (Figure 3) shows a clear positive trend across the full temperature range, with hyperthermophilic organisms clustered at high CvP values and high OGTs. The relationship is approximately linear over the mesophile and thermophile range but flattens somewhat within the hyperthermophile group, suggesting a plateau or ceiling effect at extreme temperatures.

### 3.6 PCA confirms CvP-bias as a primary axis of between-organism variation

PCA of the standardized three-metric matrix (n = 1,911 genomes with complete data) reveals that PC1 explains 47.0% of the total variance, PC2 explains 31.5%, and PC3 the remaining 21.6% (Table 3; Figure 2). The loading vector for CvP-bias on PC1 is +0.650, comparable in magnitude to the S* loading (+0.672); the mean pI loading on PC1 is +0.356. On PC2, mean pI dominates with a loading of +0.929. Thus, the three metrics are approximately orthogonal in principal component space, consistent with them measuring distinct aspects of genome composition.

**Table 3.** PCA loadings and variance explained (modern dataset, n = 1,911).

| Feature | PC1 (47.0%) | PC2 (31.5%) | PC3 (21.6%) |
| --- | --- | --- | --- |
| CvP-bias | +0.650 | −0.325 | −0.687 |
| Mean pI | +0.356 | +0.929 | −0.103 |
| $S^*$ | +0.672 | −0.178 | +0.719 |

**Figure 2.**
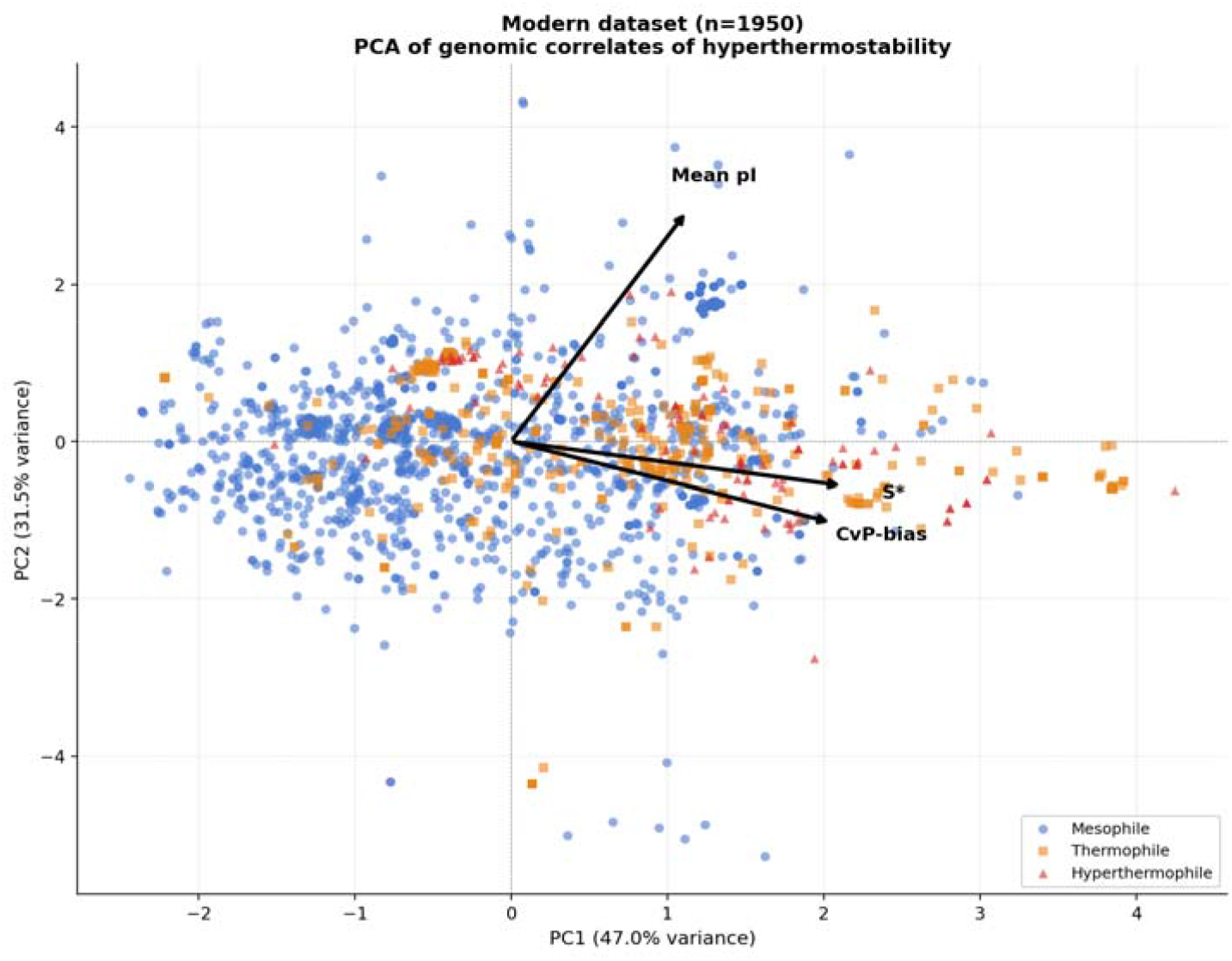
PCA biplot of 1,911 microbial genomes on the three standardized genomic metrics. Points are colored by temperature class. Loading vectors (arrows) show the direction and relative magnitude of each feature in principal component space. Hyperthermophilic organisms (red triangles) are displaced along PC1 relative to mesophiles, primarily reflecting their elevated CvP-bias.

**Figure 3.**
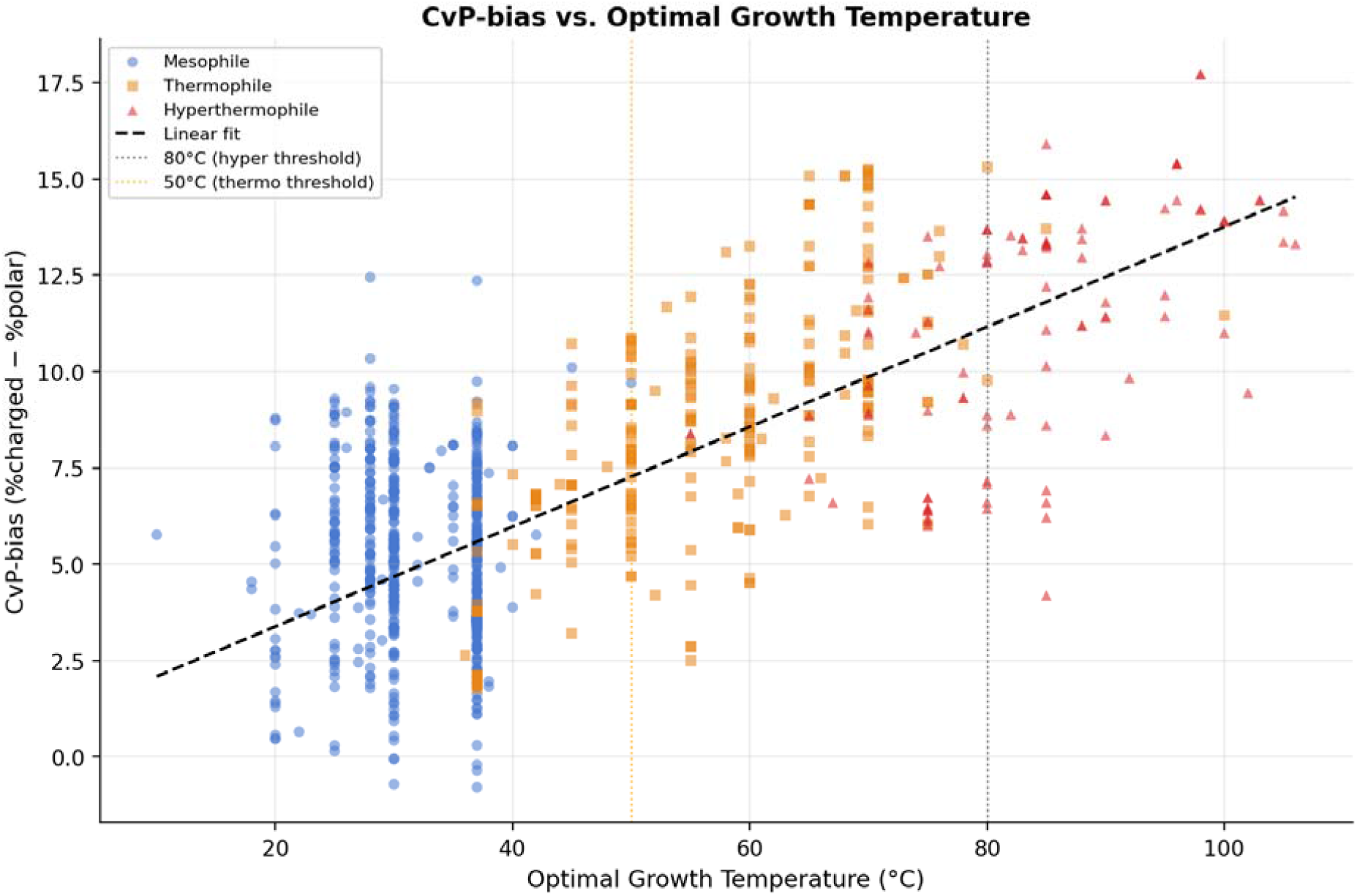
CvP-bias plotted against optimal growth temperature (OGT) for the 1,060 genomes with numeric OGT records. The dashed line shows a linear regression fit across all organisms (*r* = 0.691, *p* < 10^−150^). Vertical dotted lines indicate the conventional boundaries of thermophily (50°C) and hyperthermophily (80°C).

In the biplot (Figure 2), hyperthermophilic organisms form a cluster displaced from mesophiles and thermophiles primarily along PC1. The separation along PC1 is driven by the high CvP-bias of hyperthermophiles — despite S* loading comparably on PC1, hyperthermophiles have lower S* than mesophiles (see Section 3.3), so the opposing contributions partially cancel; the net separation along PC1 is dominated by CvP-bias. This is consistent with the 2003 finding: CvP-bias is the primary axis of discrimination between hyperthermophiles and other organisms.

## 4. Discussion

The principal finding of this study is that the CvP-bias signal identified by Suhre and Claverie in 71 genomes in 2003 [5] is confirmed, with high statistical confidence, in a dataset 28-fold larger and far more taxonomically diverse. The effect size is large (Cohen’s *d* = 2.32 for hyperthermophile vs. mesophile CvP comparison), and a single threshold on CvP-bias achieves an AUC-ROC of 0.94 for binary classification, indicating near-complete separation of temperature classes. These figures would be difficult to explain by sampling bias, taxonomic clustering, or other confounds that could in principle inflate the apparent signal in a small dataset.

### Physical interpretation

The CvP-bias reflects the relative over-representation of amino acids capable of forming salt bridges and ionic interactions (Asp, Glu, Lys, Arg, His) at the expense of polar uncharged residues that compete for surface area without the same stabilizing interactions (Asn, Gln, Ser, Thr). At elevated temperatures, hydrophobic interactions weaken, placing greater selective pressure on electrostatic and hydrogen-bonding networks to maintain folded protein structure [10,11]. The convergent evolution of high CvP-bias in phylogenetically distant hyperthermophilic Bacteria (e.g., *Aquifex, Thermotoga*) and Archaea (e.g., *Pyrococcus, Methanocaldococcus, Sulfolobus*) is consistent with independent adaptation to a shared thermal constraint.

### Mean pI

The elevation of mean proteome pI in hyperthermophiles (*r*(pI, OGT) positive but weak) is consistent with earlier reports [12] and may reflect the co-occurrence of large archaeal chromosomes in hyperthermophiles, which often encode few acidic housekeeping proteins but many basic DNA-binding and ribosomal proteins. The effect is real but modest, consistent with its secondary role in the PCA.

**Dinucleotide index S*.** The unexpected finding that hyperthermophiles have slightly *lower* S* than mesophiles — rather than higher, as one might anticipate if higher nucleotide-pair order encoded structural advantages — is also present in the original 2003 data, though it was not prominently discussed. This may reflect the predominantly AT-rich or GC-extreme genomes of many hyperthermophilic Archaea, which reduce dinucleotide diversity rather than increasing it. The S* signal, while statistically significant in ANOVA, is the weakest and least interpretable of the three metrics in this context.

### PCA interpretation

The 2003 analysis found CvP-bias to be the sole dominant loader on the first principal component, which cleanly separated hyperthermophiles from all other organisms. In the present larger dataset, PC1 loads both CvP-bias (+0.650) and S* (+0.672) with roughly equal magnitude, while PC2 is dominated by pI (+0.929). The class separation in biplot space remains clear and is driven primarily by CvP-bias, because — despite the equal loading — hyperthermophiles differ from mesophiles much more strongly on CvP-bias than on S* (standardized difference approximately +2.8 vs. −0.24, respectively). The divergence from the 2003 result may reflect the greater variance in S* captured by the larger, more diverse dataset rather than a change in the underlying biology. In both cases, the key interpretive conclusion holds: CvP-bias is the primary axis along which hyperthermophiles are separated from mesophiles.

### Limitations

The OGT metadata from BV-BRC derives from compiled literature values and may be inconsistent in precision and source. A substantial fraction of genomes (946 of 1,963) lacked numeric OGT values, limiting the regression analysis to a subset. Additionally, the dataset is taxonomically unbalanced: mesophiles are dominated by well-studied clinical and industrial organisms (Pseudomonadota, Bacillota, Actinomycetota), which may not be representative of all mesophilic diversity. The stratified sampling by phylum used here partially mitigates this imbalance but does not fully address it.

### Automated analysis by Claude Code

The entire computational workflow — from data query and bulk download to metric computation and figure generation — was designed and implemented through an interactive session with Claude Code (Anthropic, claude-sonnet-4-6), a large language model-based programming assistant operating in an agentic framework. The assistant wrote all seven Python scripts totaling approximately 900 lines of code, diagnosed and resolved API format issues, and managed parallel download of 1.65 TB of compressed genome data. The original analysis required weeks of manual effort in 2003; the present update was completed in a single automated session. This illustrates a broader methodological point: hypothesis-driven reanalysis of published findings at modern scale is becoming substantially more accessible through AI-assisted scientific programming, provided the underlying biological question and statistical approach are correctly specified by the investigator. All scripts, results tables, and figures are provided as Supplementary Data.

## 5. Conclusions

The CvP-bias — the proteome-wide difference between the fraction of charged amino acids (Asp, Glu, Lys, Arg, His) and the fraction of polar uncharged amino acids (Asn, Gln, Ser, Thr) — is confirmed as a strong, robust genomic correlate of hyperthermostability across 1,963 microbial genomes. The association is large in effect size, highly significant statistically, correlates continuously with optimal growth temperature (*r* = 0.69), and generalizes across the full taxonomic diversity of sequenced Bacteria and Archaea. These results validate the original 2003 finding [5] at modern scale, and suggest that CvP-bias can serve as a reliable, computationally inexpensive predictor of growth temperature range from proteome sequence alone.

## Acknowledgements

The author thanks Jean-Michel Claverie (Université Aix-Marseille) for the original collaboration that motivated this update. This analysis was performed using Claude Code (Anthropic) for workflow automation, BioPython for pI computations, scikit-learn for PCA, and public databases maintained by NCBI and BV-BRC.

## Data sources

*BV-BRC (bv-brc*.*org), NCBI RefSeq (ncbi*.*nlm*.*nih*.*gov/refseq/). Analysis date: April 2026. Scripts and results are provided as Supplementary Data*.

## References

1. Stetter KO. (2006) History of discovery of the first hyperthermophiles. Extremophiles 10:357–362. PMID: 16820987

2. Woese CR, Kandler O, Wheelis ML. (1990) Towards a natural system of organisms: proposal for the domains Archaea, Bacteria, and Eucarya. Proc Natl Acad Sci USA 87:4576–4579. PMID: 2112744

3. Cambillau C, Claverie JM. (2000) Structural and genomic correlates of hyperthermostability. J Biol Chem 275:32383–32386. PMID: 10938079

4. Kreil DP, Ouzounis CA. (2001) Identification of thermophilic species by the amino acid compositions deduced from their genomes. Nucleic Acids Res 29:1608–1615. PMID: 11266563

5. Suhre K, Claverie JM. (2003) Genomic correlates of hyperthermostability, an update. J Biol Chem 278:17198–17202. PMID: 12600994

6. Karlin S, Mrazek J. (1997) Compositional differences within and between eukaryotic genomes. Proc Natl Acad Sci USA 94:10227–10232. PMID: 9294192

7. Olson RD, Assaf R, Brettin T, et al. (2023) Introducing the Bacterial and Viral Bioinformatics Resource Center (BV-BRC): a resource combining PATRIC, IRD and ViPR. Nucleic Acids Res 51:D678–D689. PMID: 36350631

8. Cock PJA, Antao T, Chang JT, et al. (2009) Biopython: freely available Python tools for computational molecular biology and bioinformatics. Bioinformatics 25:1422–1423. PMID: 19304878

9. Pedregosa F, Varoquaux G, Gramfort A, et al. (2011) Scikit-learn: machine learning in Python. J Mach Learn Res 12:2825–2830.

10. Vieille C, Zeikus GJ. (2001) Hyperthermophilic enzymes: sources, uses, and molecular mechanisms for thermostability. Microbiol Mol Biol Rev 65:1–43. PMID: 11238984

11. Szilagyi A, Zavodszky P. (2000) Structural differences between mesophilic, moderately thermophilic and extremely thermophilic protein subunits: results of a comprehensive survey. Structure 8:493–504. PMID: 10801491

12. Kawashima T, Amano N, Koike H, et al. (2000) Archaeal adaptation to higher temperatures revealed by genomic sequence of Thermoplasma volcanium. Proc Natl Acad Sci USA 97:14257–14262. PMID: 11121031

13. Zeldovich KB, Berezovsky IN, Shakhnovich EI. (2007) Protein and DNA sequence determinants of thermophilic adaptation. PLoS Comput Biol 3:e5. PMID: 17222055

14. Basak S, Banerjee T, Maiti S. (2022) A positive correlation between GC content and growth temperature in prokaryotes. BMC Genomics 23:141. PMID: 35139824

15. Sauer DB, Wang DN. (2019) Predicting the optimal growth temperatures of prokaryotes using only genome derived features. Bioinformatics 35:3224–3231. PMID: 30689741

16. Sabath N, Ferrada E, Barve A, Wagner A. (2013) Growth temperature and genome size in bacteria are negatively correlated, suggesting genomic streamlining during thermal adaptation. Genome Biol Evol 5:966–977. PMID: 23563968

17. Kimura H, Mori K, Yamanaka T, Ishibashi J. (2015) Low temperature adaptation is not the opposite process of high temperature adaptation in terms of changes in amino acid composition. Genome Biol Evol 7:3437–3449. PMID: 26614525

18. Gado JE, Bhayani A. (2019) Machine learning applied to predicting microorganism growth temperatures and enzyme catalytic optima. ACS Synth Biol 8:1411–1420. PMID: 31117361

19. Ngo ST, et al. (2025) Machine learning for optimal growth temperature prediction of prokaryotes using amino acid descriptors. bioRxiv 2025.03.03.640802.

20. Nishida S, et al. (2026) Genomic and evolutionary factors influencing the prediction accuracy of optimal growth temperature in prokaryotes. bioRxiv 2025.05.30.656958. PMID: 41930963

