## Supplementary material for "Genomic correlates of hyperthermostability revisited: a large-scale validation across 1,963 microbial genomes": table1.html

### Table 1 – Genomic correlates of hyperthermostability (Suhre & Claverie 2003, reproduced)

| Organism | Class | OGT (°C) | CvP-bias | Mean pI | Dinucl S\* | # Proteins |
| --- | --- | --- | --- | --- | --- | --- |
| *Pyrococcus furiosus DSM 3638* | Hyperthermophile | 100 | +13.91 | 7.351 | 0.1697 | 2002 |
| *Pyrococcus horikoshii OT3* | Hyperthermophile | 98 | +14.21 | 7.497 | 0.1738 | 1856 |
| *Methanopyrus kandleri AV19* | Hyperthermophile | 98 | +17.72 | 6.235 | 0.1530 | 1778 |
| *Aquifex aeolicus VF5* | Hyperthermophile | 96 | +15.40 | 7.399 | 0.1960 | 1738 |
| *Pyrococcus abyssi GE5* | Hyperthermophile | 96 | +14.45 | 7.354 | 0.1577 | 1903 |
| *Aeropyrum pernix K1* | Hyperthermophile | 95 | +11.43 | 7.431 | 0.1492 | 1704 |
| *Methanocaldococcus jannaschii DSM 2661* | Hyperthermophile | 85 | +14.59 | 7.226 | 0.2057 | 1811 |
| *Archaeoglobus fulgidus DSM 4304* | Hyperthermophile | 83 | +13.46 | 6.661 | 0.1502 | 2461 |
| *Thermotoga maritima MSB8* | Hyperthermophile | 80 | +12.82 | 6.925 | 0.1783 | 1857 |
| *Sulfolobus solfataricus P2* | Hyperthermophile | 80 | +6.61 | 7.552 | 0.1128 | 2826 |
| *Thermoanaerobacter tengcongensis MB4* | Thermophile | 75 | +11.30 | 7.108 | 0.1725 | 2510 |
| *Methanothermobacter thermautotrophicus ΔH* | Thermophile | 65 | +11.33 | 6.133 | 0.1794 | 1776 |
| *Thermus thermophilus HB27* | Thermophile | 65 | +15.08 | 7.454 | 0.2429 | 2183 |
| *Thermoplasma volcanium GSS1* | Thermophile | 60 | +5.89 | 7.094 | 0.0853 | 1584 |
| *Picrophilus torridus DSM 9790* | Thermophile | 60 | +4.50 | 7.130 | 0.1689 | 1578 |
| *Geobacillus kaustophilus HTA426* | Thermophile | 60 | +9.65 | 7.032 | 0.2322 | 3369 |
| *Thermoplasma acidophilum DSM 1728* | Thermophile | 59 | +5.95 | 6.888 | 0.1152 | 1539 |
| *Thermosynechococcus elongatus BP-1* | Thermophile | 55 | +2.86 | 6.986 | 0.1593 | 2436 |
| *Escherichia coli K-12 MG1655* | Mesophile | 37 | +3.62 | 6.910 | 0.1528 | 4124 |
| *Haemophilus influenzae Rd KW20* | Mesophile | 37 | +3.80 | 6.931 | 0.1635 | 1597 |
| *Bacillus subtilis subsp. subtilis 168* | Mesophile | 37 | +6.39 | 6.849 | 0.1508 | 4251 |
| *Mycobacterium tuberculosis H37Rv* | Mesophile | 37 | +5.06 | 6.907 | 0.1451 | 3906 |
| *Mycoplasma genitalium G37* | Mesophile | 37 | +0.29 | 8.710 | 0.1760 | 504 |
| *Mycoplasma pneumoniae M129* | Mesophile | 37 | +1.25 | 8.359 | 0.1475 | 678 |
| *Chlamydia trachomatis D/UW-3/CX* | Mesophile | 37 | +3.10 | 7.165 | 0.1391 | 887 |
| *Chlamydophila pneumoniae CWL029* | Mesophile | 37 | +2.88 | 7.129 | 0.1395 | 1036 |
| *Treponema pallidum subsp. pallidum SS14* | Mesophile | 37 | +6.43 | 7.957 | 0.1178 | 976 |
| *Borrelia burgdorferi B31* | Mesophile | 37 | +5.86 | 7.997 | 0.1894 | 1376 |
| *Rickettsia prowazekii Madrid E* | Mesophile | 37 | +2.53 | 8.017 | 0.1048 | 792 |
| *Helicobacter pylori 26695* | Mesophile | 37 | +5.60 | 7.757 | 0.1946 | 1426 |
| *Neisseria meningitidis MC58* | Mesophile | 37 | +5.95 | 7.176 | 0.2124 | 1938 |
| *Pseudomonas aeruginosa PAO1* | Mesophile | 37 | +7.50 | 6.917 | 0.1588 | 5649 |
| *Campylobacter jejuni NCTC 11168* | Mesophile | 37 | +6.66 | 7.382 | 0.2041 | 1598 |
| *Halobacterium sp. NRC-1* | Mesophile | 37 | +9.22 | 5.043 | 0.2079 | 2438 |
| *Methanosarcina acetivorans C2A* | Mesophile | 37 | +6.76 | 6.446 | 0.1463 | 4627 |
| *Methanosarcina mazei Go1* | Mesophile | 37 | +7.85 | 6.424 | 0.1515 | 3341 |
| *Ureaplasma parvum serovar 3 str. ATCC 700970* | Mesophile | 37 | +2.25 | 8.129 | 0.1513 | 608 |
| *Vibrio cholerae O1 biovar El Tor str. N16961* | Mesophile | 37 | +2.97 | 6.587 | 0.1418 | 3504 |
| *Pasteurella multocida subsp. multocida Pm70* | Mesophile | 37 | +3.34 | 7.068 | 0.1438 | 2026 |
| *Brucella melitensis bv. 1 str. 16M* | Mesophile | 37 | +7.40 | 7.155 | 0.2242 | 2979 |
| *Brucella suis 1330* | Mesophile | 37 | +7.21 | 7.124 | 0.2242 | 3023 |
| *Buchnera aphidicola str. APS* | Mesophile | 37 | +3.39 | 9.008 | 0.1121 | 577 |
| *Salmonella enterica subsp. enterica serovar Typhi str. CT18* | Mesophile | 37 | +3.80 | 7.029 | 0.1591 | 4625 |
| *Yersinia pestis CO92* | Mesophile | 37 | +1.91 | 6.919 | 0.1253 | 3834 |
| *Clostridium acetobutylicum ATCC 824* | Mesophile | 37 | +5.76 | 7.443 | 0.1442 | 3835 |
| *Clostridium perfringens str. 13* | Mesophile | 37 | +8.44 | 6.785 | 0.1795 | 2656 |
| *Staphylococcus aureus subsp. aureus MW2* | Mesophile | 37 | +3.96 | 6.918 | 0.1136 | 2646 |
| *Listeria monocytogenes EGD-e* | Mesophile | 37 | +5.48 | 6.366 | 0.1274 | 2858 |
| *Listeria innocua Clip11262* | Mesophile | 37 | +5.78 | 6.458 | 0.1245 | 3057 |
| *Streptococcus pyogenes M1 GAS* | Mesophile | 37 | +4.51 | 6.861 | 0.1220 | 1701 |
| *Streptococcus pneumoniae R6* | Mesophile | 37 | +5.62 | 6.502 | 0.1246 | 1855 |
| *Enterococcus faecalis V583* | Mesophile | 37 | +3.75 | 6.631 | 0.1259 | 3165 |
| *Fusobacterium nucleatum subsp. nucleatum ATCC 25586* | Mesophile | 37 | +8.32 | 7.226 | 0.1977 | 1952 |
| *Mycobacterium leprae TN* | Mesophile | 37 | +4.59 | 6.868 | 0.1096 | 2228 |
| *Tropheryma whipplei TW08/27* | Mesophile | 37 | +2.89 | 7.909 | 0.0955 | 836 |
| *Synechocystis sp. PCC 6803* | Mesophile | 30 | +1.36 | 6.423 | 0.1876 | 3571 |
| *Caulobacter crescentus CB15* | Mesophile | 30 | +7.70 | 7.239 | 0.1783 | 3808 |
| *Sinorhizobium meliloti 1021* | Mesophile | 30 | +7.73 | 6.859 | 0.2115 | 5945 |
| *Ralstonia solanacearum GMI1000* | Mesophile | 30 | +5.34 | 7.446 | 0.1900 | 4922 |
| *Lactococcus lactis subsp. lactis Il1403* | Mesophile | 30 | +3.88 | 6.818 | 0.1506 | 2230 |
| *Deinococcus radiodurans R1* | Mesophile | 30 | +5.48 | 6.958 | 0.1567 | 3109 |
| *Corynebacterium glutamicum ATCC 13032* | Mesophile | 30 | +4.35 | 5.988 | 0.1285 | 2900 |
| *Chromobacterium violaceum ATCC 12472* | Mesophile | 30 | +6.27 | 7.228 | 0.1892 | 4332 |
| *Oceanobacillus iheyensis HTE831* | Mesophile | 30 | +4.92 | 6.269 | 0.0851 | 3488 |
| *Leptospira interrogans serovar Lai str. 56601* | Mesophile | 30 | +4.60 | 7.599 | 0.1953 | 3619 |
| *Agrobacterium fabrum str. C58* | Mesophile | 28 | +6.47 | 6.800 | 0.2117 | 5173 |
| *Mesorhizobium japonicum MAFF 303099* | Mesophile | 28 | +6.94 | 7.085 | 0.2083 | 7148 |
| *Xylella fastidiosa 9a5c* | Mesophile | 28 | +4.27 | 7.419 | 0.1351 | 2331 |
| *Xanthomonas campestris pv. campestris ATCC 33913* | Mesophile | 28 | +5.12 | 7.261 | 0.1933 | 4327 |
| *Xanthomonas oryzae pv. oryzae KACC10331* | Mesophile | 28 | +5.52 | 7.479 | 0.1919 | 3642 |
| *Nitrosomonas europaea ATCC 19718* | Mesophile | 28 | +4.84 | 6.888 | 0.1381 | 2513 |
