## Supplementary material for "Genomic correlates of hyperthermostability revisited: a large-scale validation across 1,963 microbial genomes": 01_organism_list.py

```
"""
01_organism_list.py

Build the curated list of ~71 microbial organisms from Suhre & Claverie 2003,
query NCBI for RefSeq assembly accessions and FTP paths, and write organisms.json.

Organism classification:
  hyper : T_opt > 80°C
  thermo: 50 <= T_opt <= 80°C
  meso  : T_opt < 50°C
"""

import json, time, urllib.request, urllib.parse, re, sys
from pathlib import Path

OUTFILE = Path(__file__).parent / "data" / "organisms.json"
OUTFILE.parent.mkdir(parents=True, exist_ok=True)

ENTREZ_BASE = "https://eutils.ncbi.nlm.nih.gov/entrez/eutils/"
EMAIL = ""

# ---------------------------------------------------------------------------
# Curated organism list  (name, ogt_degC, class)
# Reconstructed from: (a) original Cambillau & Claverie 2000, (b) known fully
# sequenced microbial genomes available by early 2002, (c) NCBI genome list.
# ---------------------------------------------------------------------------
ORGANISMS = [
    # ── HYPERTHERMOPHILES (T_opt > 80°C) ──────────────────────────────────
    ("Aquifex aeolicus VF5",                    96, "hyper"),
    ("Thermotoga maritima MSB8",                80, "hyper"),
    ("Pyrococcus horikoshii OT3",               98, "hyper"),
    ("Pyrococcus abyssi GE5",                   96, "hyper"),
    ("Pyrococcus furiosus DSM 3638",           100, "hyper"),
    ("Methanocaldococcus jannaschii DSM 2661",  85, "hyper"),
    ("Archaeoglobus fulgidus DSM 4304",         83, "hyper"),
    ("Aeropyrum pernix K1",                     95, "hyper"),
    ("Sulfolobus solfataricus P2",              80, "hyper"),

    # ── THERMOPHILES (50–80°C) ─────────────────────────────────────────────
    ("Thermoplasma acidophilum DSM 1728",       59, "thermo"),
    ("Thermoplasma volcanium GSS1",             60, "thermo"),
    ("Methanothermobacter thermautotrophicus ΔH", 65, "thermo"),
    ("Thermosynechococcus elongatus BP-1",      55, "thermo"),
    ("Thermoanaerobacter tengcongensis MB4",    75, "thermo"),
    ("Thermus thermophilus HB27",               65, "thermo"),
    ("Methanopyrus kandleri AV19",              98, "hyper"),   # re-classified to hyper
    ("Picrophilus torridus DSM 9790",           60, "thermo"),
    ("Geobacillus kaustophilus HTA426",         60, "thermo"),

    # ── MESOPHILES (T_opt < 50°C) ──────────────────────────────────────────
    ("Escherichia coli K-12 MG1655",            37, "meso"),
    ("Haemophilus influenzae Rd KW20",          37, "meso"),
    ("Bacillus subtilis subsp. subtilis 168",   37, "meso"),
    ("Mycobacterium tuberculosis H37Rv",        37, "meso"),
    ("Synechocystis sp. PCC 6803",              30, "meso"),
    ("Mycoplasma genitalium G37",               37, "meso"),
    ("Mycoplasma pneumoniae M129",              37, "meso"),
    ("Chlamydia trachomatis D/UW-3/CX",        37, "meso"),
    ("Chlamydophila pneumoniae CWL029",         37, "meso"),
    ("Treponema pallidum subsp. pallidum SS14", 37, "meso"),
    ("Borrelia burgdorferi B31",                37, "meso"),
    ("Rickettsia prowazekii Madrid E",          37, "meso"),
    ("Helicobacter pylori 26695",               37, "meso"),
    ("Helicobacter pylori J99",                 37, "meso"),
    ("Neisseria meningitidis MC58",             37, "meso"),
    ("Neisseria meningitidis Z2491",            37, "meso"),
    ("Pseudomonas aeruginosa PAO1",             37, "meso"),
    ("Campylobacter jejuni NCTC 11168",         37, "meso"),
    ("Halobacterium sp. NRC-1",                 37, "meso"),
    ("Methanosarcina acetivorans C2A",          37, "meso"),
    ("Methanosarcina mazei Go1",                37, "meso"),
    ("Methanobacterium thermoautotrophicum ΔH", 65, "thermo"),  # alias - will dedup
    ("Ureaplasma parvum serovar 3 str. ATCC 700970", 37, "meso"),
    ("Vibrio cholerae O1 biovar El Tor str. N16961", 37, "meso"),
    ("Pasteurella multocida subsp. multocida Pm70", 37, "meso"),
    ("Caulobacter crescentus CB15",             30, "meso"),
    ("Sinorhizobium meliloti 1021",             30, "meso"),
    ("Agrobacterium fabrum str. C58",           28, "meso"),
    ("Mesorhizobium japonicum MAFF 303099",     28, "meso"),
    ("Brucella melitensis bv. 1 str. 16M",      37, "meso"),
    ("Brucella suis 1330",                      37, "meso"),
    ("Ralstonia solanacearum GMI1000",          30, "meso"),
    ("Buchnera aphidicola str. APS",            37, "meso"),
    ("Xylella fastidiosa 9a5c",                 28, "meso"),
    ("Xanthomonas campestris pv. campestris ATCC 33913", 28, "meso"),
    ("Xanthomonas oryzae pv. oryzae KACC10331", 28, "meso"),
    ("Salmonella enterica subsp. enterica serovar Typhi str. CT18", 37, "meso"),
    ("Yersinia pestis CO92",                    37, "meso"),
    ("Clostridium acetobutylicum ATCC 824",     37, "meso"),
    ("Clostridium perfringens str. 13",         37, "meso"),
    ("Staphylococcus aureus subsp. aureus MW2", 37, "meso"),
    ("Listeria monocytogenes EGD-e",            37, "meso"),
    ("Listeria innocua Clip11262",              37, "meso"),
    ("Streptococcus pyogenes M1 GAS",           37, "meso"),
    ("Streptococcus pneumoniae R6",             37, "meso"),
    ("Lactococcus lactis subsp. lactis Il1403", 30, "meso"),
    ("Enterococcus faecalis V583",              37, "meso"),
    ("Deinococcus radiodurans R1",              30, "meso"),
    ("Fusobacterium nucleatum subsp. nucleatum ATCC 25586", 37, "meso"),
    ("Mycobacterium leprae TN",                 37, "meso"),
    ("Corynebacterium glutamicum ATCC 13032",   30, "meso"),
    ("Tropheryma whipplei TW08/27",             37, "meso"),
    ("Nitrosomonas europaea ATCC 19718",        28, "meso"),
    ("Chromobacterium violaceum ATCC 12472",    30, "meso"),
    ("Oceanobacillus iheyensis HTE831",         30, "meso"),
    ("Leptospira interrogans serovar Lai str. 56601", 30, "meso"),
]

# Deduplicate (remove Methanobacterium alias)
seen = set()
ORGANISMS_DEDUP = []
for o in ORGANISMS:
    key = o[0].split()[0:2]
    k = " ".join(key)
    if k not in seen:
        seen.add(k)
        ORGANISMS_DEDUP.append(o)

print(f"Total organisms: {len(ORGANISMS_DEDUP)}")
classes = {"hyper": 0, "thermo": 0, "meso": 0}
for _, _, c in ORGANISMS_DEDUP:
    classes[c] += 1
print(f"  hyper={classes['hyper']}  thermo={classes['thermo']}  meso={classes['meso']}")


# ---------------------------------------------------------------------------
# Query NCBI Assembly for each organism → get FTP path
# ---------------------------------------------------------------------------
def ncbi_get(endpoint, params):
    params["email"] = EMAIL
    params["tool"] = "hypertherm_repro"
    url = ENTREZ_BASE + endpoint + "?" + urllib.parse.urlencode(params)
    for attempt in range(3):
        try:
            with urllib.request.urlopen(url, timeout=30) as r:
                return json.loads(r.read())
        except Exception as e:
            if attempt == 2:
                raise
            time.sleep(2)


def find_assembly(organism_name):
    """Return (accession, ftp_path) for best RefSeq assembly, or (None,None)."""
    # Try RefSeq reference first, then representative, then any complete
    for filter_str in [
        f'"{organism_name}"[ORGN] AND "reference genome"[PROP] AND "Complete Genome"[ASMSTAT]',
        f'"{organism_name}"[ORGN] AND "representative genome"[PROP]',
        f'"{organism_name}"[ORGN] AND "Complete Genome"[ASMSTAT]',
        f'"{organism_name}"[ORGN] AND "Chromosome"[ASMSTAT]',
    ]:
        data = ncbi_get("esearch.fcgi", {"db": "assembly", "term": filter_str,
                                          "retmax": 5, "retmode": "json"})
        ids = data.get("esearchresult", {}).get("idlist", [])
        if ids:
            # Fetch summary to get FTP path
            sum_data = ncbi_get("esummary.fcgi", {"db": "assembly",
                                                   "id": ",".join(ids[:3]),
                                                   "retmode": "json"})
            for uid, rec in sum_data.get("result", {}).items():
                if uid == "uids":
                    continue
                ftp = rec.get("ftppath_refseq", "") or rec.get("ftppath_genbank", "")
                acc = rec.get("assemblyaccession", "")
                if ftp and ftp != "na":
                    return acc, ftp
        time.sleep(0.34)  # NCBI rate limit: max 3/sec
    return None, None


print("\nQuerying NCBI Assembly for FTP paths ...")
results = []
for i, (name, ogt, cls) in enumerate(ORGANISMS_DEDUP):
    acc, ftp = find_assembly(name)
    status = "✓" if acc else "✗"
    print(f"  [{i+1:3d}/{len(ORGANISMS_DEDUP)}] {status} {name[:60]:<60}  {acc or 'NOT FOUND'}")
    results.append({
        "name": name,
        "ogt": ogt,
        "class": cls,
        "accession": acc,
        "ftp_path": ftp,
    })
    time.sleep(0.34)

found = sum(1 for r in results if r["accession"])
print(f"\nFound FTP paths: {found}/{len(results)}")

with open(OUTFILE, "w") as f:
    json.dump(results, f, indent=2)
print(f"Written: {OUTFILE}")
```
