## Supplementary material for "Genomic correlates of hyperthermostability revisited: a large-scale validation across 1,963 microbial genomes": 02_download_data.py

```
"""
02_download_data.py

Download protein FASTA (*_protein.faa.gz) and CDS nucleotide FASTA
(*_cds_from_genomic.fna.gz) for each organism in data/organisms.json.

Saves files to data/raw/{sanitized_name}/
Uses checkpointing: skips organisms already downloaded.
Rate: ~3 requests/sec (NCBI FTP policy).
"""

import json, gzip, time, urllib.request, urllib.error, re, sys
from pathlib import Path

BASE_DIR  = Path(__file__).parent
ORG_FILE  = BASE_DIR / "data" / "organisms.json"
RAW_DIR   = BASE_DIR / "data" / "raw"
RAW_DIR.mkdir(parents=True, exist_ok=True)

TIMEOUT = 120  # seconds per file

def sanitize(name):
    """Turn organism name into a safe directory name."""
    return re.sub(r'[^\w\-]', '_', name)[:80]

def ftp_to_https(ftp_url):
    """Convert ftp:// URL to https:// for more reliable access."""
    return ftp_url.replace("ftp://ftp.ncbi.nlm.nih.gov", "https://ftp.ncbi.nlm.nih.gov")

def list_ftp_dir(ftp_path):
    """Get directory listing via HTTPS to find file names."""
    url = ftp_to_https(ftp_path) + "/"
    try:
        with urllib.request.urlopen(url, timeout=30) as r:
            html = r.read().decode()
        # Extract filenames from directory listing
        files = re.findall(r'href="([^"]+)"', html)
        return [f for f in files if not f.startswith('/') and f != '../']
    except Exception as e:
        print(f"    [list_dir error] {e}")
        return []

def download_file(url, dest_path):
    """Download url → dest_path. Returns True on success."""
    dest_path = Path(dest_path)
    if dest_path.exists() and dest_path.stat().st_size > 0:
        return True  # already downloaded
    dest_path.parent.mkdir(parents=True, exist_ok=True)
    try:
        with urllib.request.urlopen(url, timeout=TIMEOUT) as r, \
             open(dest_path, 'wb') as f:
            while chunk := r.read(1 << 20):
                f.write(chunk)
        return True
    except Exception as e:
        print(f"    [download error] {url}: {e}")
        if dest_path.exists():
            dest_path.unlink()
        return False

def get_genome_files(ftp_path, org_dir):
    """
    Find and download protein.faa.gz and cds_from_genomic.fna.gz
    from an NCBI FTP assembly directory.
    Returns (protein_path, cds_path) or (None, None) on failure.
    """
    files = list_ftp_dir(ftp_path)
    base_url = ftp_to_https(ftp_path)

    protein_file = None
    cds_file     = None

    for f in files:
        if f.endswith("_protein.faa.gz"):
            protein_file = f
        elif f.endswith("_cds_from_genomic.fna.gz"):
            cds_file = f

    results = {}
    for label, fname in [("protein", protein_file), ("cds", cds_file)]:
        if fname:
            url  = f"{base_url}/{fname}"
            dest = org_dir / fname
            ok   = download_file(url, dest)
            results[label] = dest if ok else None
        else:
            results[label] = None

    return results.get("protein"), results.get("cds")


# ── Main ────────────────────────────────────────────────────────────────────
with open(ORG_FILE) as f:
    organisms = json.load(f)

done = skipped = failed = 0
checkpoint_file = BASE_DIR / "data" / "download_status.json"
if checkpoint_file.exists():
    with open(checkpoint_file) as f:
        checkpoint = json.load(f)
else:
    checkpoint = {}

for i, org in enumerate(organisms):
    name = org["name"]
    safe = sanitize(name)
    ftp  = org.get("ftp_path")

    if not ftp:
        print(f"[{i+1:2d}/{len(organisms)}] SKIP (no FTP): {name[:60]}")
        failed += 1
        continue

    org_dir = RAW_DIR / safe

    # Check checkpoint
    if safe in checkpoint and checkpoint[safe].get("status") == "ok":
        prot_path = checkpoint[safe].get("protein")
        cds_path  = checkpoint[safe].get("cds")
        if prot_path and Path(prot_path).exists():
            print(f"[{i+1:2d}/{len(organisms)}] CACHED: {name[:60]}")
            skipped += 1
            continue

    print(f"[{i+1:2d}/{len(organisms)}] Downloading: {name[:60]}")
    org_dir.mkdir(parents=True, exist_ok=True)

    prot_path, cds_path = get_genome_files(ftp, org_dir)

    if prot_path:
        # Quick validation: decompress first bytes
        try:
            with gzip.open(prot_path, 'rt') as gz:
                first = gz.read(100)
            if not first.startswith('>'):
                raise ValueError("Not a FASTA file")
            print(f"    ✓ protein: {prot_path.name}  ({prot_path.stat().st_size // 1024} KB)")
        except Exception as e:
            print(f"    ✗ protein validation failed: {e}")
            prot_path.unlink(missing_ok=True)
            prot_path = None

    if cds_path:
        try:
            with gzip.open(cds_path, 'rt') as gz:
                first = gz.read(100)
            if not first.startswith('>'):
                raise ValueError("Not a FASTA file")
            print(f"    ✓ CDS:     {cds_path.name}  ({cds_path.stat().st_size // 1024} KB)")
        except Exception as e:
            print(f"    ✗ CDS validation failed: {e}")
            cds_path.unlink(missing_ok=True)
            cds_path = None

    status = "ok" if prot_path else "no_protein"
    checkpoint[safe] = {
        "status":  status,
        "protein": str(prot_path) if prot_path else None,
        "cds":     str(cds_path)  if cds_path  else None,
    }
    with open(checkpoint_file, 'w') as f:
        json.dump(checkpoint, f, indent=2)

    if prot_path:
        done += 1
    else:
        failed += 1

    time.sleep(0.4)  # be polite to NCBI

print(f"\nDone: {done} downloaded, {skipped} cached, {failed} failed")
```
