## Supplementary material for "Genomic correlates of hyperthermostability revisited: a large-scale validation across 1,963 microbial genomes": 03_compute_metrics.py

```
"""
03_compute_metrics.py

Compute three metrics for each organism:
  1. CvP-bias   = %{D,E,K,R,H} − %{N,Q,S,T}     (protein FASTA)
  2. mean_pI    = average isoelectric point         (protein FASTA)
  3. dinucl_S   = dinucleotide deviation index S*  (CDS FASTA)
                  S* = (1/16) × Σ|ρ*(XY) − 1|
                  ρ*(XY) = f(XY) / (f(X)·f(Y))

Output: results/metrics_original.csv
"""

import gzip, json, re, sys
from collections import Counter
from pathlib import Path

import numpy as np
import pandas as pd
from Bio.SeqUtils.ProtParam import ProteinAnalysis

BASE_DIR  = Path(__file__).parent
ORG_FILE  = BASE_DIR / "data" / "organisms.json"
CHK_FILE  = BASE_DIR / "data" / "download_status.json"
OUT_FILE  = BASE_DIR / "results" / "metrics_original.csv"
OUT_FILE.parent.mkdir(parents=True, exist_ok=True)

CHARGED = set("DEKRH")
POLAR   = set("NQST")
AMBIG   = set("BJOUXZ*-")   # skip these in AA counting


# ── Amino-acid composition helpers ─────────────────────────────────────────
def iter_fasta(gz_path):
    """Yield (header, seq) from a .faa.gz file."""
    header, parts = None, []
    with gzip.open(gz_path, 'rt') as f:
        for line in f:
            line = line.rstrip()
            if line.startswith('>'):
                if header is not None:
                    yield header, ''.join(parts)
                header, parts = line[1:], []
            else:
                parts.append(line.upper())
    if header is not None:
        yield header, ''.join(parts)


def compute_cvp(seq):
    """CvP-bias for a single (concatenated) protein sequence."""
    counts = Counter(seq)
    total  = sum(counts[aa] for aa in counts if aa not in AMBIG)
    if total == 0:
        return np.nan
    c = sum(counts[aa] for aa in CHARGED) / total * 100
    p = sum(counts[aa] for aa in POLAR)   / total * 100
    return c - p


def compute_mean_pI(faa_gz):
    """Average isoelectric point over all proteins in a proteome."""
    pis = []
    for _, seq in iter_fasta(faa_gz):
        # Strip ambiguous/stop characters
        clean = re.sub(r'[^ACDEFGHIKLMNPQRSTVWY]', '', seq)
        if len(clean) < 3:
            continue
        try:
            pi = ProteinAnalysis(clean).isoelectric_point()
            pis.append(pi)
        except Exception:
            pass
    return float(np.mean(pis)) if pis else np.nan


# ── Dinucleotide deviation index ────────────────────────────────────────────
NUCLEOTIDES = list("ACGT")
DINUCLEOTIDES = [a + b for a in NUCLEOTIDES for b in NUCLEOTIDES]

def compute_dinucl_S(cds_gz):
    """
    S* = (1/16) × Σ|ρ*(XY) − 1|
    ρ*(XY) = f(XY) / (f(X)·f(Y))
    Computed on pooled coding sequences.
    """
    mono  = Counter()
    di    = Counter()

    with gzip.open(cds_gz, 'rt') as f:
        for line in f:
            line = line.rstrip()
            if line.startswith('>'):
                continue
            seq = line.upper()
            # count mono
            for nt in seq:
                if nt in "ACGT":
                    mono[nt] += 1
            # count di (only canonical)
            for i in range(len(seq) - 1):
                a, b = seq[i], seq[i+1]
                if a in "ACGT" and b in "ACGT":
                    di[a+b] += 1

    total_mono = sum(mono.values())
    total_di   = sum(di.values())
    if total_mono == 0 or total_di == 0:
        return np.nan

    f_mono = {nt: mono[nt] / total_mono for nt in "ACGT"}
    f_di   = {dn: di[dn] / total_di     for dn in DINUCLEOTIDES}

    deviations = []
    for dn in DINUCLEOTIDES:
        expected = f_mono[dn[0]] * f_mono[dn[1]]
        if expected > 0:
            rho = f_di[dn] / expected
            deviations.append(abs(rho - 1.0))

    return float(np.mean(deviations)) if deviations else np.nan


# ── Main ────────────────────────────────────────────────────────────────────
with open(ORG_FILE)  as f: organisms = json.load(f)
with open(CHK_FILE)  as f: checkpoint = json.load(f)

# Map safe name → paths
def sanitize(name):
    return re.sub(r'[^\w\-]', '_', name)[:80]

rows = []
for org in organisms:
    name  = org["name"]
    safe  = sanitize(name)
    ogt   = org["ogt"]
    cls   = org["class"]

    info  = checkpoint.get(safe, {})
    prot  = info.get("protein")
    cds   = info.get("cds")

    print(f"Processing: {name[:55]:<55}", end="  ")

    if not prot or not Path(prot).exists():
        print("SKIP (no protein file)")
        continue

    # Concatenate all AA sequences for CvP (fast path)
    all_aa = []
    for _, seq in iter_fasta(prot):
        all_aa.append(seq)
    concat = ''.join(all_aa)

    cvp    = compute_cvp(concat)
    mean_pi = compute_mean_pI(prot)

    dinucl = np.nan
    if cds and Path(cds).exists():
        dinucl = compute_dinucl_S(cds)

    print(f"CvP={cvp:+.2f}  pI={mean_pi:.3f}  S*={dinucl:.4f}" if not np.isnan(dinucl)
          else f"CvP={cvp:+.2f}  pI={mean_pi:.3f}  S*=NA")

    rows.append({
        "organism": name,
        "class":    cls,
        "ogt":      ogt,
        "cvp":      round(cvp,    4),
        "mean_pI":  round(mean_pi, 4),
        "dinucl_S": round(dinucl,  6) if not np.isnan(dinucl) else np.nan,
        "n_proteins": len(all_aa),
    })

df = pd.DataFrame(rows)
df.to_csv(OUT_FILE, index=False)
print(f"\nSaved {len(df)} rows → {OUT_FILE}")

# Quick summary
print("\n── Mean metrics by class ──")
summary = df.groupby("class")[["cvp","mean_pI","dinucl_S"]].agg(["mean","std"]).round(3)
print(summary)
```
