## Supplementary material for "Genomic correlates of hyperthermostability revisited: a large-scale validation across 1,963 microbial genomes": 04_reproduce_figures.py

```
"""
04_reproduce_figures.py

Reproduce Table 1 and Figures 1 & 2 from Suhre & Claverie 2003.

Figure 1 — Distribution of three metrics by organism class
Figure 2 — PCA biplot showing CvP dominates discrimination
Table 1  — Sorted organism table (CSV + HTML)
"""

import numpy as np
import pandas as pd
import matplotlib
matplotlib.use('Agg')
import matplotlib.pyplot as plt
import matplotlib.patches as mpatches
from matplotlib.ticker import MultipleLocator
from sklearn.preprocessing import StandardScaler
from sklearn.decomposition import PCA
from pathlib import Path

BASE_DIR = Path(__file__).parent
METRICS  = BASE_DIR / "results" / "metrics_original.csv"
OUT_DIR  = BASE_DIR / "results"
OUT_DIR.mkdir(exist_ok=True)

# ── Colors and labels ────────────────────────────────────────────────────────
CLASS_COLOR  = {"meso": "#4878CF", "thermo": "#E8861A", "hyper": "#D62728"}
CLASS_LABEL  = {"meso": "Mesophile", "thermo": "Thermophile", "hyper": "Hyperthermophile"}
CLASS_ORDER  = ["meso", "thermo", "hyper"]
CLASS_MARKER = {"meso": "o", "thermo": "s", "hyper": "^"}

df = pd.read_csv(METRICS)
df["class_label"] = df["class"].map(CLASS_LABEL)

# ── Table 1 ──────────────────────────────────────────────────────────────────
# Sort: hypers first, then thermos, then mesos; within each class by OGT desc
order_map = {"hyper": 0, "thermo": 1, "meso": 2}
df_sorted = df.assign(sort_order=df["class"].map(order_map)) \
              .sort_values(["sort_order", "ogt"], ascending=[True, False]) \
              .drop(columns="sort_order")

table1 = df_sorted[["organism","class","ogt","cvp","mean_pI","dinucl_S","n_proteins"]].copy()
table1.columns = ["Organism", "Class", "OGT (°C)", "CvP-bias", "Mean pI", "Dinucl S*", "# Proteins"]

table1.to_csv(OUT_DIR / "table1.csv", index=False)

# HTML version with color-coded rows
def row_style(cls):
    bg = {"hyper":"#FFD5D5","thermo":"#FFE8CC","meso":"#D5E8FF"}
    return f'background-color:{bg[cls]}'

html_rows = []
for _, r in df_sorted.iterrows():
    style = row_style(r["class"])
    html_rows.append(
        f'<tr style="{style}">'
        f'<td><i>{r["organism"]}</i></td>'
        f'<td>{CLASS_LABEL[r["class"]]}</td>'
        f'<td style="text-align:right">{r["ogt"]}</td>'
        f'<td style="text-align:right">{r["cvp"]:+.2f}</td>'
        f'<td style="text-align:right">{r["mean_pI"]:.3f}</td>'
        f'<td style="text-align:right">{r["dinucl_S"]:.4f}</td>'
        f'<td style="text-align:right">{r["n_proteins"]}</td>'
        f'</tr>'
    )

html = f"""<!DOCTYPE html><html><head>
<style>
  body {{font-family:Arial,sans-serif; font-size:12px}}
  table {{border-collapse:collapse; width:100%}}
  th {{background:#333; color:white; padding:6px 10px; text-align:left}}
  td {{padding:4px 10px; border-bottom:1px solid #ddd}}
</style></head><body>
<h2>Table 1 – Genomic correlates of hyperthermostability (Suhre &amp; Claverie 2003, reproduced)</h2>
<table>
<tr><th>Organism</th><th>Class</th><th>OGT (°C)</th>
    <th>CvP-bias</th><th>Mean pI</th><th>Dinucl S*</th><th># Proteins</th></tr>
{''.join(html_rows)}
</table></body></html>"""

with open(OUT_DIR / "table1.html", "w") as f:
    f.write(html)

print(f"Table 1 written ({len(df)} organisms)")

# ── Figure 1 — Metric distributions ─────────────────────────────────────────
fig, axes = plt.subplots(1, 3, figsize=(14, 5))
fig.suptitle("Genomic correlates of hyperthermostability\n(Suhre & Claverie 2003, reproduced)",
             fontsize=13, fontweight='bold', y=1.01)

metrics_info = [
    ("cvp",      "CvP-bias (%charged − %polar)",   "CvP-bias (%)"),
    ("mean_pI",  "Average proteome pI",              "Mean pI"),
    ("dinucl_S", "Dinucleotide deviation index S*",  "S*"),
]

np.random.seed(42)
for ax, (col, title, ylabel) in zip(axes, metrics_info):
    for cls in CLASS_ORDER:
        vals = df[df["class"] == cls][col].dropna().values
        x_pos = CLASS_ORDER.index(cls)
        color = CLASS_COLOR[cls]

        # Box plot
        bp = ax.boxplot(vals, positions=[x_pos], widths=0.45,
                        patch_artist=True, notch=False, zorder=2,
                        boxprops=dict(facecolor=color, alpha=0.35, linewidth=1.5),
                        medianprops=dict(color="black", linewidth=2),
                        whiskerprops=dict(linewidth=1.5),
                        capprops=dict(linewidth=1.5),
                        flierprops=dict(marker='', markersize=0))

        # Jitter strip
        jitter = (np.random.rand(len(vals)) - 0.5) * 0.25
        ax.scatter(x_pos + jitter, vals, color=color, s=40, alpha=0.85,
                   edgecolors='white', linewidths=0.5, zorder=3,
                   marker=CLASS_MARKER[cls])

    ax.set_xticks(range(3))
    ax.set_xticklabels([CLASS_LABEL[c] for c in CLASS_ORDER], fontsize=9)
    ax.set_ylabel(ylabel, fontsize=10)
    ax.set_title(title, fontsize=10, pad=6)
    ax.spines[['top','right']].set_visible(False)
    ax.grid(axis='y', alpha=0.3)

    # Annotate N
    for i, cls in enumerate(CLASS_ORDER):
        n = (df["class"] == cls).sum()
        ax.text(i, ax.get_ylim()[0], f"n={n}", ha='center', va='bottom',
                fontsize=8, color='gray')

# Legend
patches = [mpatches.Patch(color=CLASS_COLOR[c], label=CLASS_LABEL[c])
           for c in CLASS_ORDER]
fig.legend(handles=patches, loc='upper right', framealpha=0.9,
           bbox_to_anchor=(1.0, 1.0), fontsize=9)

plt.tight_layout()
fig.savefig(OUT_DIR / "figure1_original.png", dpi=200, bbox_inches='tight')
plt.close()
print("Figure 1 saved")


# ── Figure 2 — PCA biplot ────────────────────────────────────────────────────
feat_cols  = ["cvp", "mean_pI", "dinucl_S"]
feat_names = ["CvP-bias", "Mean pI", "S*"]

df_pca = df[feat_cols + ["class", "organism", "ogt"]].dropna()
X = df_pca[feat_cols].values
scaler = StandardScaler()
X_scaled = scaler.fit_transform(X)

pca = PCA(n_components=2)
scores = pca.fit_transform(X_scaled)
loadings = pca.components_  # shape (2, 3)

var_exp = pca.explained_variance_ratio_ * 100
print(f"\nPCA variance explained: PC1={var_exp[0]:.1f}%  PC2={var_exp[1]:.1f}%")
print("Loadings (PC1, PC2):")
for name, l1, l2 in zip(feat_names, loadings[0], loadings[1]):
    print(f"  {name:<12}  PC1={l1:+.3f}  PC2={l2:+.3f}")

fig, ax = plt.subplots(figsize=(9, 7))

# Scatter organisms
for _, row in df_pca.iterrows():
    idx = df_pca.index.get_loc(_)
    sc_x, sc_y = scores[idx]
    cls = row["class"]
    ax.scatter(sc_x, sc_y,
               color=CLASS_COLOR[cls],
               marker=CLASS_MARKER[cls],
               s=70, alpha=0.8,
               edgecolors='white', linewidths=0.5, zorder=3)

# Annotate hyperthermophiles
for _, row in df_pca[df_pca["class"] == "hyper"].iterrows():
    idx = df_pca.index.get_loc(_)
    sc_x, sc_y = scores[idx]
    short = row["organism"].split()[0] + " " + row["organism"].split()[1]
    ax.annotate(short, (sc_x, sc_y), fontsize=7,
                xytext=(4, 3), textcoords='offset points', color='#8B0000')

# Loading vectors (biplot arrows)
scale = 2.0  # scale factor for visibility
for name, l1, l2 in zip(feat_names, loadings[0], loadings[1]):
    ax.annotate("", xy=(l1*scale, l2*scale), xytext=(0, 0),
                arrowprops=dict(arrowstyle='->', color='black', lw=2))
    ax.text(l1*scale*1.12, l2*scale*1.12, name, fontsize=10, fontweight='bold',
            ha='center', va='center', color='black')

ax.axhline(0, color='gray', lw=0.5, ls='--')
ax.axvline(0, color='gray', lw=0.5, ls='--')
ax.set_xlabel(f"PC1 ({var_exp[0]:.1f}% variance)", fontsize=11)
ax.set_ylabel(f"PC2 ({var_exp[1]:.1f}% variance)", fontsize=11)
ax.set_title("PCA of genomic correlates of hyperthermostability\n(Suhre & Claverie 2003, reproduced)",
             fontsize=12, fontweight='bold')

patches = [mpatches.Patch(color=CLASS_COLOR[c], label=CLASS_LABEL[c])
           for c in CLASS_ORDER]
ax.legend(handles=patches, loc='lower right', framealpha=0.9, fontsize=9)
ax.spines[['top','right']].set_visible(False)
ax.grid(alpha=0.25)

plt.tight_layout()
fig.savefig(OUT_DIR / "figure2_original.png", dpi=200, bbox_inches='tight')
plt.close()
print("Figure 2 saved")

# ── Summary stats ────────────────────────────────────────────────────────────
print("\n── Per-class statistics ──")
for cls in CLASS_ORDER:
    sub = df[df["class"] == cls]
    print(f"\n{CLASS_LABEL[cls]} (n={len(sub)}):")
    for col in ["cvp","mean_pI","dinucl_S"]:
        v = sub[col].dropna()
        print(f"  {col:<12}  mean={v.mean():+.3f}  std={v.std():.3f}  "
              f"min={v.min():+.3f}  max={v.max():+.3f}")
```
