## Supplementary material for "Genomic correlates of hyperthermostability revisited: a large-scale validation across 1,963 microbial genomes": 05_modern_query.py

```
"""
05_modern_query.py

1. Query BV-BRC for all bacterial/archaeal genomes with temperature annotations.
2. Download NCBI RefSeq assembly_summary.txt for bacteria + archaea to get
   FTP paths in bulk (one file lookup vs thousands of API calls).
3. Match BV-BRC genomes to NCBI assembly FTP paths.
4. Save final list to data/modern_genomes.csv.

Total available: ~9,000 genomes with temperature annotations.
We select all hypers + thermos, and up to 1,500 mesophiles.
"""

import json, time, urllib.request, gzip, io, re
from pathlib import Path
from collections import Counter

import pandas as pd
import numpy as np

BASE_DIR = Path(__file__).parent
DATA_DIR = BASE_DIR / "data"
OUT_FILE = DATA_DIR / "modern_genomes.csv"
DATA_DIR.mkdir(parents=True, exist_ok=True)

# ── Step 1: Fetch BV-BRC genome metadata ────────────────────────────────────
print("=== Step 1: Fetching BV-BRC genome metadata ===")

SELECT_FIELDS = ",".join([
    "genome_id","genome_name","taxon_id","superkingdom",
    "phylum","class","order","family","genus","species",
    "optimal_temperature","temperature_range",
    "genome_quality","reference_genome","genome_status",
    "assembly_accession","ncbi_project_id",
])

TR_VALUES = ["Mesophilic","Thermophilic","Hyperthermophilic","Psychrophilic"]

all_records = []
for tr in TR_VALUES:
    print(f"  Querying {tr} ...")
    offset, batch = 0, 5000
    while True:
        url = (f"https://www.bv-brc.org/api/genome/"
               f"?eq(temperature_range,{tr})"
               f"&select({SELECT_FIELDS})"
               f"&limit({batch},{offset})")
        req = urllib.request.Request(url, headers={"Accept":"application/json"})
        try:
            with urllib.request.urlopen(req, timeout=60) as r:
                records = json.loads(r.read())
            if not records:
                break
            all_records.extend(records)
            print(f"    {tr}: {offset + len(records)} records fetched")
            if len(records) < batch:
                break
            offset += batch
            time.sleep(0.3)
        except Exception as e:
            print(f"    Error at offset {offset}: {e}")
            break

print(f"Total BV-BRC records: {len(all_records)}")

df_bvbrc = pd.DataFrame(all_records)

# Classify thermo class
def classify(row):
    tr = str(row.get("temperature_range","")).lower()
    if "hyperthermophil" in tr: return "hyper"
    if "thermophil" in tr:      return "thermo"
    if "mesophil" in tr:        return "meso"
    if "psychro" in tr:         return "psychro"
    return "unknown"

df_bvbrc["thermo_class"] = df_bvbrc.apply(classify, axis=1)
df_bvbrc["optimal_temperature"] = pd.to_numeric(
    df_bvbrc["optimal_temperature"].astype(str).str.extract(r'([\d.]+)')[0],
    errors='coerce'
)

# Keep only bacteria + archaea (BV-BRC uses 'superkingdom' or 'kingdom')
king_col = "superkingdom" if "superkingdom" in df_bvbrc.columns else "kingdom"
if king_col in df_bvbrc.columns:
    df_bvbrc = df_bvbrc[df_bvbrc[king_col].isin(["Bacteria","Archaea","bacteria","archaea"])]

# Drop records without assembly_accession
df_bvbrc = df_bvbrc[df_bvbrc["assembly_accession"].notna() &
                     (df_bvbrc["assembly_accession"] != "")]
df_bvbrc = df_bvbrc[df_bvbrc["thermo_class"].isin(["meso","thermo","hyper"])]

print(f"\nAfter filtering (bacteria/archaea with assembly_accession):")
print(df_bvbrc["thermo_class"].value_counts())

# ── Step 2: Download NCBI assembly_summary.txt ───────────────────────────────
print("\n=== Step 2: Downloading NCBI assembly summary files ===")

asm_summary_paths = []
for domain in ["bacteria", "archaea"]:
    url = f"https://ftp.ncbi.nlm.nih.gov/genomes/refseq/{domain}/assembly_summary.txt"
    local = DATA_DIR / f"assembly_summary_{domain}.txt.gz"

    if local.exists():
        print(f"  Cached: assembly_summary_{domain}.txt.gz")
    else:
        print(f"  Downloading assembly_summary_{domain}.txt.gz ...")
        try:
            with urllib.request.urlopen(url, timeout=120) as r, \
                 gzip.open(local, 'wb') as f_out:
                # Re-compress as gzip for storage
                data = r.read()
                f_out.write(data)
            print(f"    Saved {local.stat().st_size // 1024} KB (uncompressed {len(data)//1024} KB)")
        except Exception as e:
            print(f"  FALLBACK: downloading without compression")
            local = DATA_DIR / f"assembly_summary_{domain}.txt"
            with urllib.request.urlopen(url, timeout=120) as r, \
                 open(local, 'wb') as f_out:
                f_out.write(r.read())
    asm_summary_paths.append(local)


# ── Step 3: Parse assembly summaries → accession→ftp_path dict ──────────────
print("\n=== Step 3: Building accession → FTP path lookup ===")

def parse_assembly_summary(path):
    """Parse NCBI assembly_summary.txt and return dict: accession → ftp_path."""
    lookup = {}
    path = Path(path)

    opener = gzip.open if str(path).endswith('.gz') else open
    with opener(path, 'rt', errors='replace') as f:
        for line in f:
            if line.startswith('#') or not line.strip():
                continue
            parts = line.rstrip('\n').split('\t')
            if len(parts) < 20:
                continue
            # Columns: 0=assembly_accession, 1=bioproject, ..., 19=ftp_path
            acc  = parts[0]  # e.g. GCF_000000000.1
            ftp  = parts[19] # ftp_path
            if ftp and ftp != 'na':
                lookup[acc] = ftp
                # Also map without version number
                acc_noversion = acc.rsplit('.',1)[0]
                if acc_noversion not in lookup:
                    lookup[acc_noversion] = ftp
    return lookup

acc_to_ftp = {}
for p in asm_summary_paths:
    lk = parse_assembly_summary(p)
    acc_to_ftp.update(lk)
    print(f"  Loaded {len(lk)} entries from {Path(p).name}")

print(f"Total lookup entries: {len(acc_to_ftp)}")


# ── Step 4: Match BV-BRC genomes to FTP paths ───────────────────────────────
print("\n=== Step 4: Matching BV-BRC genomes to NCBI FTP paths ===")

def normalize_acc(acc):
    """Normalize accession: try GCF version, GCA version, no-version."""
    if not isinstance(acc, str): return []
    candidates = [acc]
    if acc.startswith("GCA_"):
        candidates.append(acc.replace("GCA_","GCF_"))
    elif acc.startswith("GCF_"):
        candidates.append(acc.replace("GCF_","GCA_"))
    # Also try without version
    for a in list(candidates):
        candidates.append(a.rsplit('.',1)[0])
    return candidates

def find_ftp(acc):
    for cand in normalize_acc(acc):
        if cand in acc_to_ftp:
            return acc_to_ftp[cand]
    return None

df_bvbrc["ftp_path"] = df_bvbrc["assembly_accession"].apply(find_ftp)
n_matched = df_bvbrc["ftp_path"].notna().sum()
print(f"Matched {n_matched}/{len(df_bvbrc)} genomes to NCBI FTP paths")
print(f"Per class (matched only):")
print(df_bvbrc[df_bvbrc["ftp_path"].notna()]["thermo_class"].value_counts())


# ── Step 5: Select representative subset ────────────────────────────────────
print("\n=== Step 5: Selecting representative subset ===")

df_matched = df_bvbrc[df_bvbrc["ftp_path"].notna()].copy()

# All hypers + thermos; stratified sample of mesophiles
df_hyper  = df_matched[df_matched["thermo_class"] == "hyper"]
df_thermo = df_matched[df_matched["thermo_class"] == "thermo"]
df_meso   = df_matched[df_matched["thermo_class"] == "meso"]

# Sample mesophiles: aim for ~1500, stratified by phylum
N_MESO_SAMPLE = 1500
if len(df_meso) > N_MESO_SAMPLE:
    # Stratified by phylum; cap each phylum proportionally
    df_meso_sample = (df_meso.groupby("phylum", group_keys=False)
                              .apply(lambda x: x.sample(
                                  n=max(1, int(N_MESO_SAMPLE * len(x) / len(df_meso))),
                                  random_state=42
                              )))
    if len(df_meso_sample) < N_MESO_SAMPLE:
        # Top up if rounding left us short
        remaining = df_meso[~df_meso.index.isin(df_meso_sample.index)]
        extras = remaining.sample(n=min(N_MESO_SAMPLE - len(df_meso_sample), len(remaining)),
                                  random_state=42)
        df_meso_sample = pd.concat([df_meso_sample, extras])
else:
    df_meso_sample = df_meso

df_final = pd.concat([df_hyper, df_thermo, df_meso_sample]).reset_index(drop=True)
print(f"Final selection: {len(df_final)} genomes")
print(df_final["thermo_class"].value_counts())
print(f"Kingdoms: {df_final[king_col].value_counts().to_dict() if king_col in df_final.columns else 'N/A'}")

df_final.to_csv(OUT_FILE, index=False)
print(f"\nSaved → {OUT_FILE}")

# Summary statistics
print("\n── OGT distribution per class ──")
for cls in ["hyper","thermo","meso"]:
    sub = df_final[df_final["thermo_class"]==cls]["optimal_temperature"].dropna()
    print(f"  {cls:<8} n={len(df_final[df_final['thermo_class']==cls]):4d}  "
          f"OGT={sub.mean():.1f}±{sub.std():.1f}°C  "
          f"[{sub.min():.0f}–{sub.max():.0f}]" if len(sub)>0 else f"  {cls}: no OGT data")
```
