## Supplementary material for "Genomic correlates of hyperthermostability revisited: a large-scale validation across 1,963 microbial genomes": 06_modern_download.py

```
"""
06_modern_download.py

Parallel bulk download of protein FASTAs and CDS FASTAs for all 2,017
modern genomes listed in data/modern_genomes.csv.

Uses 10 parallel threads. Skips files already downloaded.
Writes download progress to data/modern_download_status.json.
"""

import json, gzip, re, time, urllib.request
from concurrent.futures import ThreadPoolExecutor, as_completed
from pathlib import Path
from threading import Lock

import pandas as pd

BASE_DIR  = Path(__file__).parent
CSV_FILE  = BASE_DIR / "data" / "modern_genomes.csv"
RAW_DIR   = BASE_DIR / "data" / "modern_raw"
CHK_FILE  = BASE_DIR / "data" / "modern_download_status.json"
RAW_DIR.mkdir(parents=True, exist_ok=True)

MAX_WORKERS = 10
TIMEOUT     = 120   # seconds per file download

# ── Helpers ──────────────────────────────────────────────────────────────────
def sanitize(name):
    return re.sub(r'[^\w\-]', '_', str(name))[:80]

def ftp_to_https(ftp_url):
    return str(ftp_url).replace("ftp://ftp.ncbi.nlm.nih.gov",
                                "https://ftp.ncbi.nlm.nih.gov")

def list_dir(ftp_path):
    """List NCBI FTP directory via HTTPS."""
    url = ftp_to_https(ftp_path).rstrip('/') + "/"
    try:
        with urllib.request.urlopen(url, timeout=30) as r:
            html = r.read().decode(errors='replace')
        return re.findall(r'href="([^"/?]+)"', html)
    except Exception:
        return []

def download_file(url, dest):
    dest = Path(dest)
    if dest.exists() and dest.stat().st_size > 100:
        return True
    dest.parent.mkdir(parents=True, exist_ok=True)
    try:
        with urllib.request.urlopen(url, timeout=TIMEOUT) as r, \
             open(dest, 'wb') as f:
            while chunk := r.read(1 << 20):
                f.write(chunk)
        return True
    except Exception as e:
        if dest.exists(): dest.unlink(missing_ok=True)
        return False

def validate_gz_fasta(path):
    """Quick check that file is a valid gzipped FASTA."""
    try:
        with gzip.open(path, 'rt') as f:
            first = f.read(100)
        return first.lstrip().startswith('>')
    except Exception:
        return False


# ── Load checkpoint ───────────────────────────────────────────────────────────
lock = Lock()
if CHK_FILE.exists():
    with open(CHK_FILE) as f:
        checkpoint = json.load(f)
else:
    checkpoint = {}

def save_checkpoint():
    with lock, open(CHK_FILE, 'w') as f:
        json.dump(checkpoint, f, indent=2)

# ── Per-genome download function ─────────────────────────────────────────────
def download_genome(row):
    """Download protein.faa.gz + cds.fna.gz for one genome. Returns status dict."""
    genome_id = str(row["genome_id"])
    name      = str(row.get("genome_name", genome_id))
    ftp       = str(row.get("ftp_path", ""))
    cls       = str(row.get("thermo_class", "unknown"))

    safe_id   = sanitize(genome_id)

    with lock:
        cached = checkpoint.get(safe_id)
    if cached and cached.get("status") == "ok":
        prot = cached.get("protein")
        if prot and Path(prot).exists():
            return {"id": genome_id, "status": "cached", "protein": prot,
                    "cds": cached.get("cds"), "class": cls, "name": name}

    if not ftp or ftp == "nan":
        return {"id": genome_id, "status": "no_ftp", "class": cls, "name": name}

    org_dir = RAW_DIR / cls / safe_id
    org_dir.mkdir(parents=True, exist_ok=True)

    files   = list_dir(ftp)
    base_url = ftp_to_https(ftp).rstrip('/')

    protein_file = next((f for f in files if f.endswith("_protein.faa.gz")), None)
    cds_file     = next((f for f in files if f.endswith("_cds_from_genomic.fna.gz")), None)

    prot_path = cds_path = None

    if protein_file:
        dest = org_dir / protein_file
        ok   = download_file(f"{base_url}/{protein_file}", dest)
        if ok and validate_gz_fasta(dest):
            prot_path = str(dest)
        elif dest.exists():
            dest.unlink(missing_ok=True)

    if cds_file:
        dest = org_dir / cds_file
        ok   = download_file(f"{base_url}/{cds_file}", dest)
        if ok and validate_gz_fasta(dest):
            cds_path = str(dest)
        elif dest.exists():
            dest.unlink(missing_ok=True)

    status = "ok" if prot_path else "no_protein"
    result = {"id": genome_id, "status": status, "protein": prot_path,
              "cds": cds_path, "class": cls, "name": name}

    with lock:
        checkpoint[safe_id] = {"status": status, "protein": prot_path, "cds": cds_path}
    save_checkpoint()

    return result


# ── Main ─────────────────────────────────────────────────────────────────────
df = pd.read_csv(CSV_FILE)
print(f"Downloading {len(df)} genomes with {MAX_WORKERS} parallel workers...")
print(f"Classes: {df['thermo_class'].value_counts().to_dict()}")

# Check what's already done
already_done = sum(
    1 for _, row in df.iterrows()
    if checkpoint.get(sanitize(str(row["genome_id"])), {}).get("status") == "ok"
    and checkpoint.get(sanitize(str(row["genome_id"])), {}).get("protein")
    and Path(checkpoint[sanitize(str(row["genome_id"]))]["protein"]).exists()
)
print(f"Already downloaded: {already_done}/{len(df)}")

rows_list = [row for _, row in df.iterrows()]

done = skipped = failed = 0
start_time = time.time()

with ThreadPoolExecutor(max_workers=MAX_WORKERS) as executor:
    futures = {executor.submit(download_genome, row): row for row in rows_list}
    for i, future in enumerate(as_completed(futures)):
        r = future.result()
        status = r["status"]
        if status == "ok":
            done += 1
            tag = "✓"
        elif status == "cached":
            skipped += 1
            tag = "~"
        else:
            failed += 1
            tag = "✗"

        elapsed = time.time() - start_time
        total_processed = done + skipped + failed
        rate = total_processed / elapsed if elapsed > 0 else 0
        eta  = (len(rows_list) - total_processed) / rate if rate > 0 else 0

        if (i + 1) % 50 == 0 or status == "ok" or total_processed <= 5:
            print(f"  [{total_processed:4d}/{len(rows_list)}] {tag} {r['name'][:45]:<45}  "
                  f"(done={done} cached={skipped} fail={failed}  "
                  f"rate={rate:.1f}/s  ETA={eta/60:.1f}min)")

save_checkpoint()
print(f"\nDone: {done} downloaded, {skipped} cached, {failed} failed")
print(f"Total time: {(time.time()-start_time)/60:.1f} min")

# Status summary
with open(CHK_FILE) as f:
    chk = json.load(f)
ok_count = sum(1 for v in chk.values() if v.get("status")=="ok")
print(f"Total successfully downloaded in checkpoint: {ok_count}")
```
