## Supplementary material for "Genomic correlates of hyperthermostability revisited: a large-scale validation across 1,963 microbial genomes": 07_modern_analysis.py

```
"""
07_modern_analysis.py

Compute CvP-bias, mean pI, and dinucleotide S* for all successfully downloaded
modern genomes. Combines with the original 71-organism results and generates
updated Table 1 equivalent and Figures 1 & 2 for the modern dataset.
"""

import gzip, json, re, sys
from collections import Counter
from pathlib import Path

import numpy as np
import pandas as pd
import matplotlib
matplotlib.use('Agg')
import matplotlib.pyplot as plt
import matplotlib.patches as mpatches
from sklearn.preprocessing import StandardScaler
from sklearn.decomposition import PCA
from Bio.SeqUtils.ProtParam import ProteinAnalysis

BASE_DIR   = Path(__file__).parent
CHK_FILE   = BASE_DIR / "data" / "modern_download_status.json"
META_FILE  = BASE_DIR / "data" / "modern_genomes.csv"
OUT_METRICS = BASE_DIR / "results" / "metrics_modern.csv"
OUT_DIR     = BASE_DIR / "results"
OUT_DIR.mkdir(exist_ok=True)

CHARGED = set("DEKRH")
POLAR   = set("NQST")
AMBIG   = set("BJOUXZ*-")

CLASS_COLOR  = {"meso": "#4878CF", "thermo": "#E8861A", "hyper": "#D62728"}
CLASS_LABEL  = {"meso": "Mesophile", "thermo": "Thermophile", "hyper": "Hyperthermophile"}
CLASS_ORDER  = ["meso", "thermo", "hyper"]
CLASS_MARKER = {"meso": "o", "thermo": "s", "hyper": "^"}


# ── Metric functions (same as original) ──────────────────────────────────────
def iter_fasta(gz_path):
    header, parts = None, []
    with gzip.open(gz_path, 'rt') as f:
        for line in f:
            line = line.rstrip()
            if line.startswith('>'):
                if header is not None:
                    yield header, ''.join(parts)
                header, parts = line[1:], []
            else:
                parts.append(line.upper())
    if header is not None:
        yield header, ''.join(parts)

def compute_cvp(concat_seq):
    counts = Counter(concat_seq)
    total  = sum(counts[aa] for aa in counts if aa not in AMBIG)
    if total == 0: return np.nan
    c = sum(counts[aa] for aa in CHARGED) / total * 100
    p = sum(counts[aa] for aa in POLAR)   / total * 100
    return c - p

def compute_mean_pI(faa_gz):
    pis = []
    for _, seq in iter_fasta(faa_gz):
        clean = re.sub(r'[^ACDEFGHIKLMNPQRSTVWY]', '', seq)
        if len(clean) < 3: continue
        try:
            pis.append(ProteinAnalysis(clean).isoelectric_point())
        except Exception:
            pass
    return float(np.mean(pis)) if pis else np.nan

NUCLEOTIDES = list("ACGT")
DINUCLEOTIDES = [a+b for a in NUCLEOTIDES for b in NUCLEOTIDES]

def compute_dinucl_S(cds_gz):
    mono, di = Counter(), Counter()
    with gzip.open(cds_gz, 'rt') as f:
        for line in f:
            line = line.rstrip()
            if line.startswith('>'): continue
            seq = line.upper()
            for nt in seq:
                if nt in "ACGT": mono[nt] += 1
            for i in range(len(seq)-1):
                a, b = seq[i], seq[i+1]
                if a in "ACGT" and b in "ACGT": di[a+b] += 1
    total_mono = sum(mono.values())
    total_di   = sum(di.values())
    if total_mono == 0 or total_di == 0: return np.nan
    f_mono = {nt: mono[nt]/total_mono for nt in "ACGT"}
    f_di   = {dn: di[dn]/total_di     for dn in DINUCLEOTIDES}
    devs   = [abs(f_di[dn]/(f_mono[dn[0]]*f_mono[dn[1]]) - 1.0)
              for dn in DINUCLEOTIDES if f_mono[dn[0]]*f_mono[dn[1]] > 0]
    return float(np.mean(devs)) if devs else np.nan


# ── Load checkpoint & metadata ───────────────────────────────────────────────
with open(CHK_FILE) as f: checkpoint = json.load(f)
df_meta = pd.read_csv(META_FILE)

# Build id → metadata map
meta_map = {str(r["genome_id"]): r for _, r in df_meta.iterrows()}

# ── Compute metrics ───────────────────────────────────────────────────────────
rows = []
ok_entries = [(k, v) for k, v in checkpoint.items()
              if v.get("status") == "ok" and v.get("protein") and Path(v["protein"]).exists()]

print(f"Computing metrics for {len(ok_entries)} downloaded genomes ...")

for i, (safe_id, info) in enumerate(ok_entries):
    prot = info["protein"]
    cds  = info.get("cds")

    # Recover genome_id from path (stored as hyper/thermo/meso/{safe_id}/filename)
    # The safe_id IS the sanitized genome_id
    # Find genome_id in metadata by matching safe_id
    genome_id = None
    for gid in meta_map:
        if re.sub(r'[^\w\-]', '_', str(gid))[:80] == safe_id:
            genome_id = gid
            break

    if genome_id is None:
        continue

    meta = meta_map[genome_id]
    cls  = str(meta.get("thermo_class", "unknown"))
    ogt  = meta.get("optimal_temperature", np.nan)
    name = str(meta.get("genome_name", genome_id))

    if (i + 1) % 100 == 0:
        print(f"  [{i+1:4d}/{len(ok_entries)}] {name[:50]}")

    try:
        all_aa = []
        for _, seq in iter_fasta(prot):
            all_aa.append(seq)
        concat = ''.join(all_aa)

        cvp     = compute_cvp(concat)
        mean_pi = compute_mean_pI(prot)
        dinucl  = compute_dinucl_S(cds) if cds and Path(cds).exists() else np.nan

        rows.append({
            "organism":   name,
            "genome_id":  genome_id,
            "class":      cls,
            "ogt":        ogt,
            "cvp":        round(cvp,    4),
            "mean_pI":    round(mean_pi, 4),
            "dinucl_S":   round(dinucl,  6) if not np.isnan(dinucl) else np.nan,
            "n_proteins": len(all_aa),
            "source":     "modern",
        })
    except Exception as e:
        print(f"  ERROR {name[:40]}: {e}")

df_modern = pd.DataFrame(rows)
df_modern.to_csv(OUT_METRICS, index=False)
print(f"\nSaved {len(df_modern)} rows → {OUT_METRICS}")

print("\n── Mean metrics by class ──")
summary = df_modern.groupby("class")[["cvp","mean_pI","dinucl_S"]].agg(["mean","std"]).round(3)
print(summary)


# ── Combine with original dataset ────────────────────────────────────────────
orig_file = BASE_DIR / "results" / "metrics_original.csv"
df_orig = pd.read_csv(orig_file)
df_orig["source"] = "original_2003"
df_modern["source"] = "modern"

df_combined = pd.concat([df_orig, df_modern], ignore_index=True)
df_combined.to_csv(OUT_DIR / "metrics_combined.csv", index=False)


# ── Figure 1 modern — Metric distributions ───────────────────────────────────
for df_plot, tag, subtitle in [
    (df_modern, "modern", f"Modern dataset (n={len(df_modern)})"),
    (df_combined, "combined", f"Combined 2003 + modern (n={len(df_combined)})"),
]:
    fig, axes = plt.subplots(1, 3, figsize=(14, 5))
    fig.suptitle(f"Genomic correlates of hyperthermostability — {subtitle}",
                 fontsize=12, fontweight='bold', y=1.01)

    metrics_info = [
        ("cvp",      "CvP-bias (%charged − %polar)",   "CvP-bias (%)"),
        ("mean_pI",  "Average proteome pI",              "Mean pI"),
        ("dinucl_S", "Dinucleotide deviation index S*",  "S*"),
    ]

    np.random.seed(42)
    for ax, (col, title, ylabel) in zip(axes, metrics_info):
        for cls in CLASS_ORDER:
            vals = df_plot[df_plot["class"] == cls][col].dropna().values
            if len(vals) == 0: continue
            x_pos = CLASS_ORDER.index(cls)
            color = CLASS_COLOR[cls]

            ax.boxplot(vals, positions=[x_pos], widths=0.5,
                       patch_artist=True, notch=False, zorder=2,
                       boxprops=dict(facecolor=color, alpha=0.3, linewidth=1.5),
                       medianprops=dict(color="black", linewidth=2.5),
                       whiskerprops=dict(linewidth=1.5),
                       capprops=dict(linewidth=1.5),
                       flierprops=dict(marker='', markersize=0))

            # Violin overlay for large N
            if len(vals) > 20:
                parts = ax.violinplot(vals, positions=[x_pos], widths=0.45,
                                      showmeans=False, showmedians=False, showextrema=False)
                for pc in parts['bodies']:
                    pc.set_facecolor(color)
                    pc.set_alpha(0.18)

            # Jitter (subsample for large datasets)
            n_show = min(len(vals), 200)
            idx    = np.random.choice(len(vals), n_show, replace=False)
            jitter = (np.random.rand(n_show) - 0.5) * 0.3
            ax.scatter(x_pos + jitter, vals[idx],
                       color=color, s=18, alpha=0.55,
                       edgecolors='none', zorder=3,
                       marker=CLASS_MARKER[cls])

        ax.set_xticks(range(3))
        ax.set_xticklabels([CLASS_LABEL[c] for c in CLASS_ORDER], fontsize=9)
        ax.set_ylabel(ylabel, fontsize=10)
        ax.set_title(title, fontsize=10, pad=6)
        ax.spines[['top','right']].set_visible(False)
        ax.grid(axis='y', alpha=0.3)
        ymin = ax.get_ylim()[0]
        for j, cls in enumerate(CLASS_ORDER):
            n = (df_plot["class"] == cls).sum()
            ax.text(j, ymin, f"n={n}", ha='center', va='bottom', fontsize=8, color='gray')

    patches = [mpatches.Patch(color=CLASS_COLOR[c], label=CLASS_LABEL[c]) for c in CLASS_ORDER]
    fig.legend(handles=patches, loc='upper right', framealpha=0.9,
               bbox_to_anchor=(1.0, 1.0), fontsize=9)
    plt.tight_layout()
    fig.savefig(OUT_DIR / f"figure1_{tag}.png", dpi=200, bbox_inches='tight')
    plt.close()
    print(f"Figure 1 ({tag}) saved")


# ── Figure 2 modern — PCA biplot ──────────────────────────────────────────────
feat_cols  = ["cvp", "mean_pI", "dinucl_S"]
feat_names = ["CvP-bias", "Mean pI", "S*"]

for df_plot, tag, subtitle in [
    (df_modern,   "modern",   f"Modern dataset (n={{n}})"),
    (df_combined, "combined", f"Combined (n={{n}})"),
]:
    df_pca = df_plot[feat_cols + ["class","organism"]].dropna()
    n = len(df_pca)
    X        = df_pca[feat_cols].values
    X_scaled = StandardScaler().fit_transform(X)
    pca      = PCA(n_components=2)
    scores   = pca.fit_transform(X_scaled)
    loadings = pca.components_
    var_exp  = pca.explained_variance_ratio_ * 100

    print(f"\nPCA ({tag}): PC1={var_exp[0]:.1f}%  PC2={var_exp[1]:.1f}%")
    print("Loadings:")
    for name, l1, l2 in zip(feat_names, loadings[0], loadings[1]):
        print(f"  {name:<12}  PC1={l1:+.3f}  PC2={l2:+.3f}")

    fig, ax = plt.subplots(figsize=(10, 8))

    for cls in CLASS_ORDER:
        mask = df_pca["class"].values == cls
        if not mask.any(): continue
        ax.scatter(scores[mask, 0], scores[mask, 1],
                   color=CLASS_COLOR[cls], marker=CLASS_MARKER[cls],
                   s=30, alpha=0.6, edgecolors='none', zorder=3, label=CLASS_LABEL[cls])

    # Biplot loading arrows
    scale = np.abs(scores).max() * 0.6
    for name, l1, l2 in zip(feat_names, loadings[0], loadings[1]):
        ax.annotate("", xy=(l1*scale, l2*scale), xytext=(0, 0),
                    arrowprops=dict(arrowstyle='->', color='black', lw=2.5))
        ax.text(l1*scale*1.15, l2*scale*1.15, name, fontsize=11, fontweight='bold',
                ha='center', va='center', color='black')

    ax.axhline(0, color='gray', lw=0.5, ls='--')
    ax.axvline(0, color='gray', lw=0.5, ls='--')
    ax.set_xlabel(f"PC1 ({var_exp[0]:.1f}% variance)", fontsize=11)
    ax.set_ylabel(f"PC2 ({var_exp[1]:.1f}% variance)", fontsize=11)
    ax.set_title(subtitle.format(n=n) + "\nPCA of genomic correlates of hyperthermostability",
                 fontsize=12, fontweight='bold')
    ax.legend(loc='lower right', framealpha=0.9, fontsize=9)
    ax.spines[['top','right']].set_visible(False)
    ax.grid(alpha=0.2)
    plt.tight_layout()
    fig.savefig(OUT_DIR / f"figure2_{tag}.png", dpi=200, bbox_inches='tight')
    plt.close()
    print(f"Figure 2 ({tag}) saved")


# ── Figure 3 — CvP-bias vs OGT scatter ───────────────────────────────────────
for df_plot, tag, subtitle in [
    (df_combined, "combined", "CvP-bias vs. Optimal Growth Temperature"),
]:
    df_ogt = df_plot[["ogt","cvp","class"]].dropna()
    if len(df_ogt) == 0: continue

    fig, ax = plt.subplots(figsize=(9, 6))
    for cls in CLASS_ORDER:
        sub = df_ogt[df_ogt["class"] == cls]
        ax.scatter(sub["ogt"], sub["cvp"],
                   color=CLASS_COLOR[cls], marker=CLASS_MARKER[cls],
                   s=25, alpha=0.55, edgecolors='none', label=CLASS_LABEL[cls])

    # Fit polynomial trendline across all
    from numpy.polynomial import polynomial as P
    x_all = df_ogt["ogt"].values
    y_all = df_ogt["cvp"].values
    coefs = P.polyfit(x_all, y_all, 1)
    x_line = np.linspace(x_all.min(), x_all.max(), 200)
    ax.plot(x_line, P.polyval(x_line, coefs), 'k--', lw=1.5, label='Linear fit')

    ax.axvline(80, color='gray', lw=1, ls=':', label='80°C (hyper threshold)')
    ax.axvline(50, color='orange', lw=1, ls=':', alpha=0.7, label='50°C (thermo threshold)')

    ax.set_xlabel("Optimal Growth Temperature (°C)", fontsize=11)
    ax.set_ylabel("CvP-bias (%charged − %polar)", fontsize=11)
    ax.set_title(subtitle, fontsize=12, fontweight='bold')
    ax.legend(loc='upper left', fontsize=8, framealpha=0.9)
    ax.spines[['top','right']].set_visible(False)
    ax.grid(alpha=0.25)
    plt.tight_layout()
    fig.savefig(OUT_DIR / f"figure3_cvp_vs_ogt_{tag}.png", dpi=200, bbox_inches='tight')
    plt.close()
    print(f"Figure 3 (CvP vs OGT) saved")

print("\nAll figures saved to results/")
```
