## Supplementary figures and images for "Genomic correlates of hyperthermostability revisited: a large-scale validation across 1,963 microbial genomes"

### figure1_combined.png

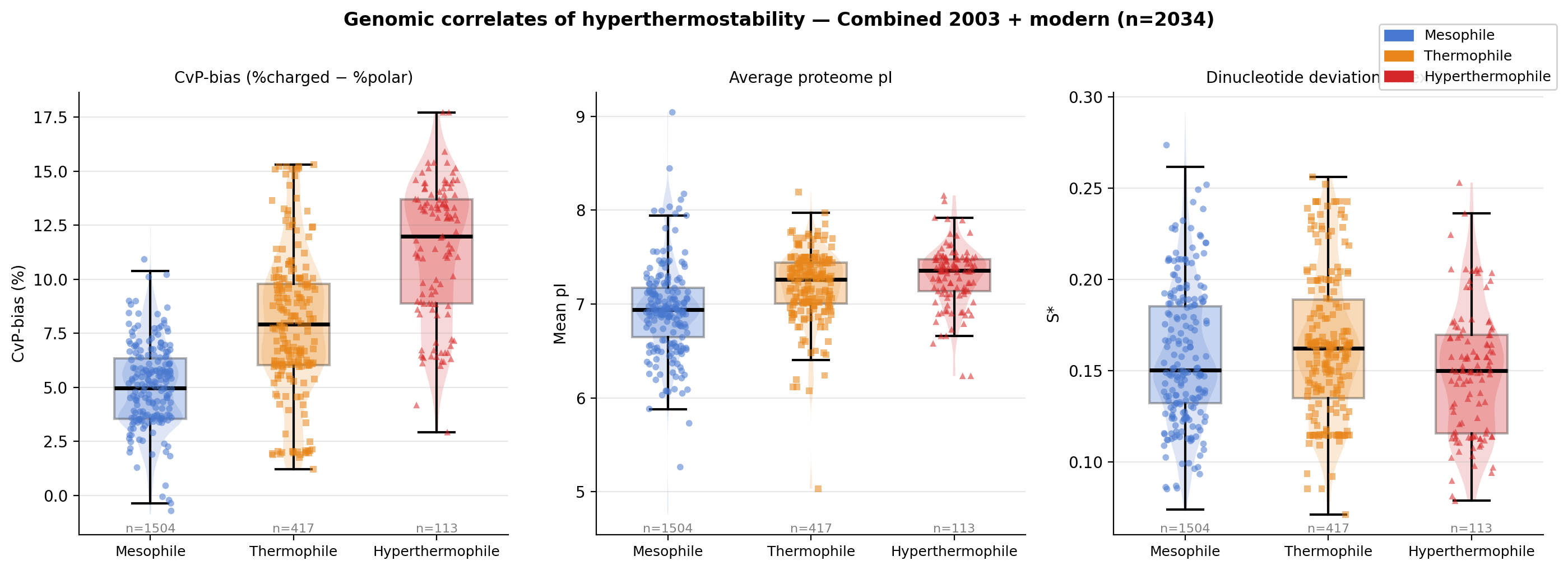

### figure1_modern.png

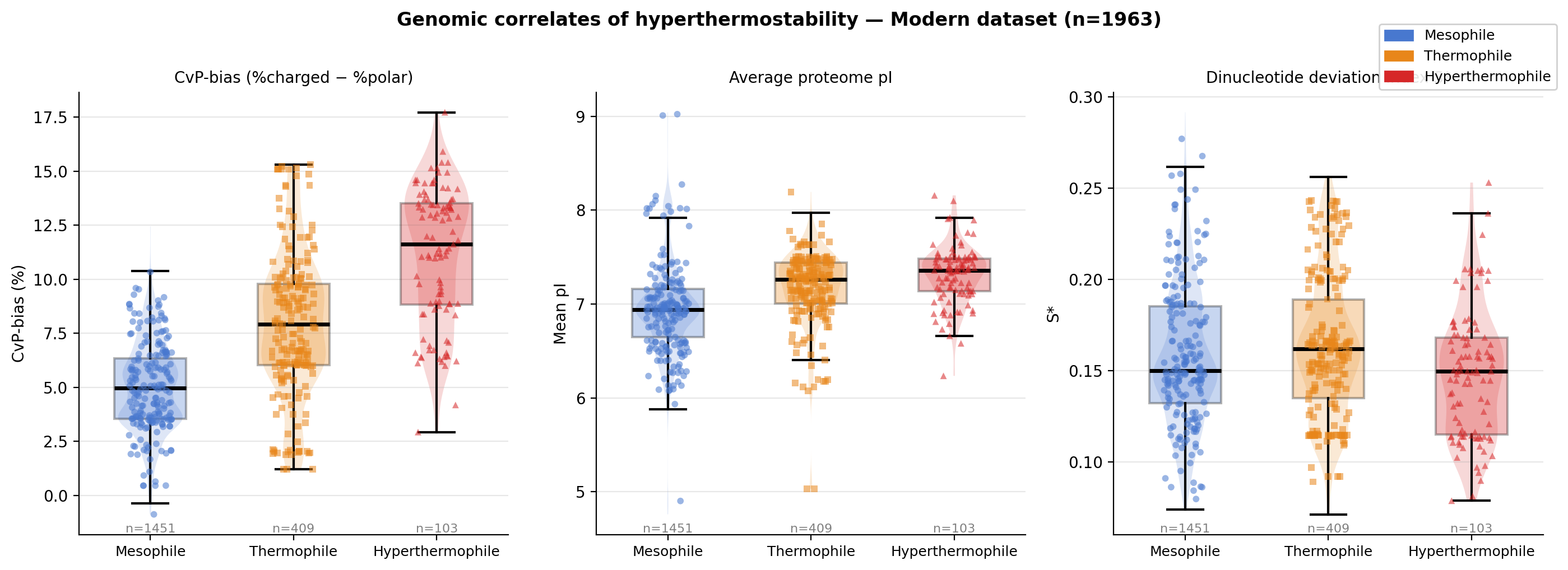

### figure1_original.png

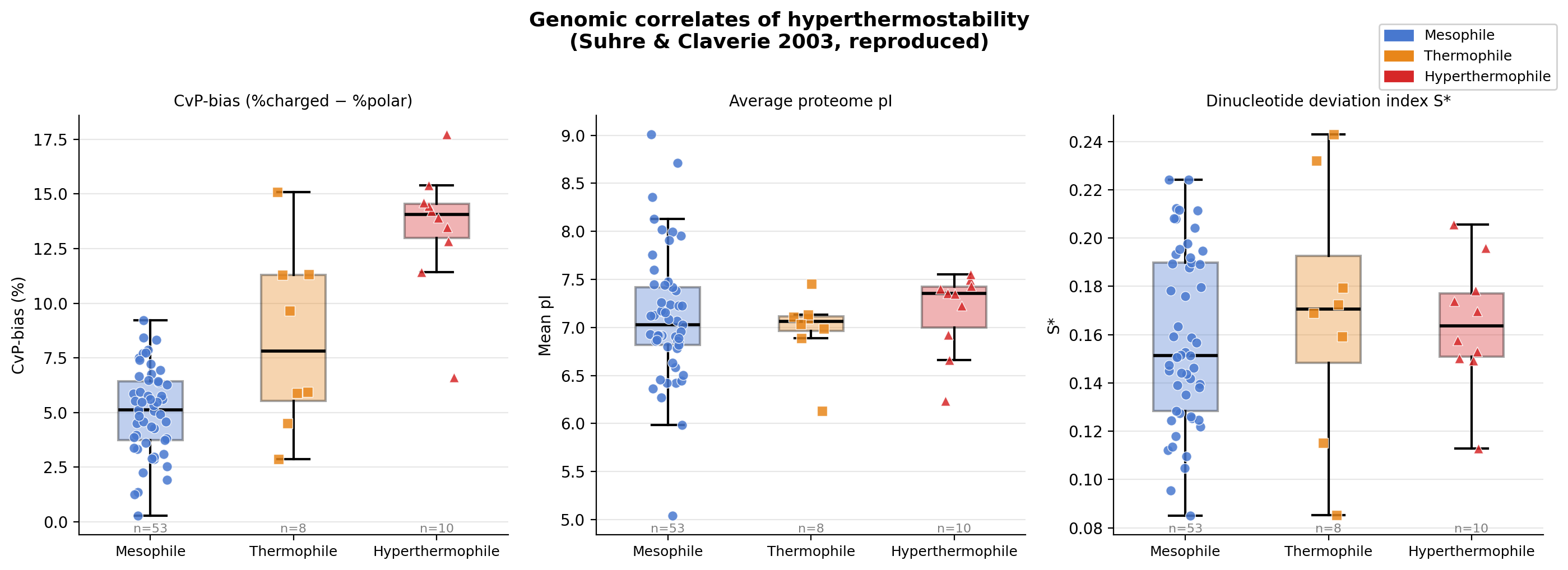

### figure2_combined.png

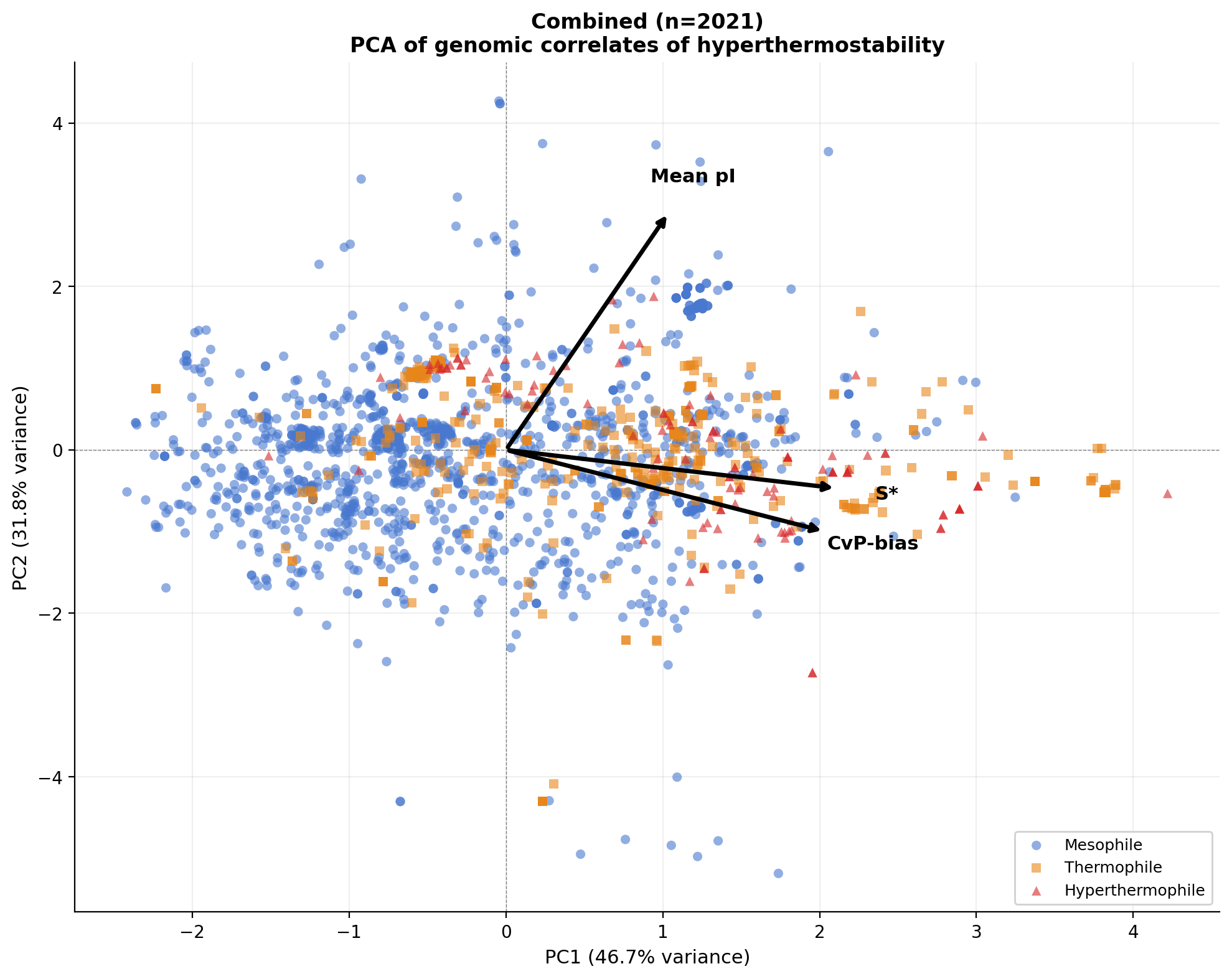

### figure2_modern.png

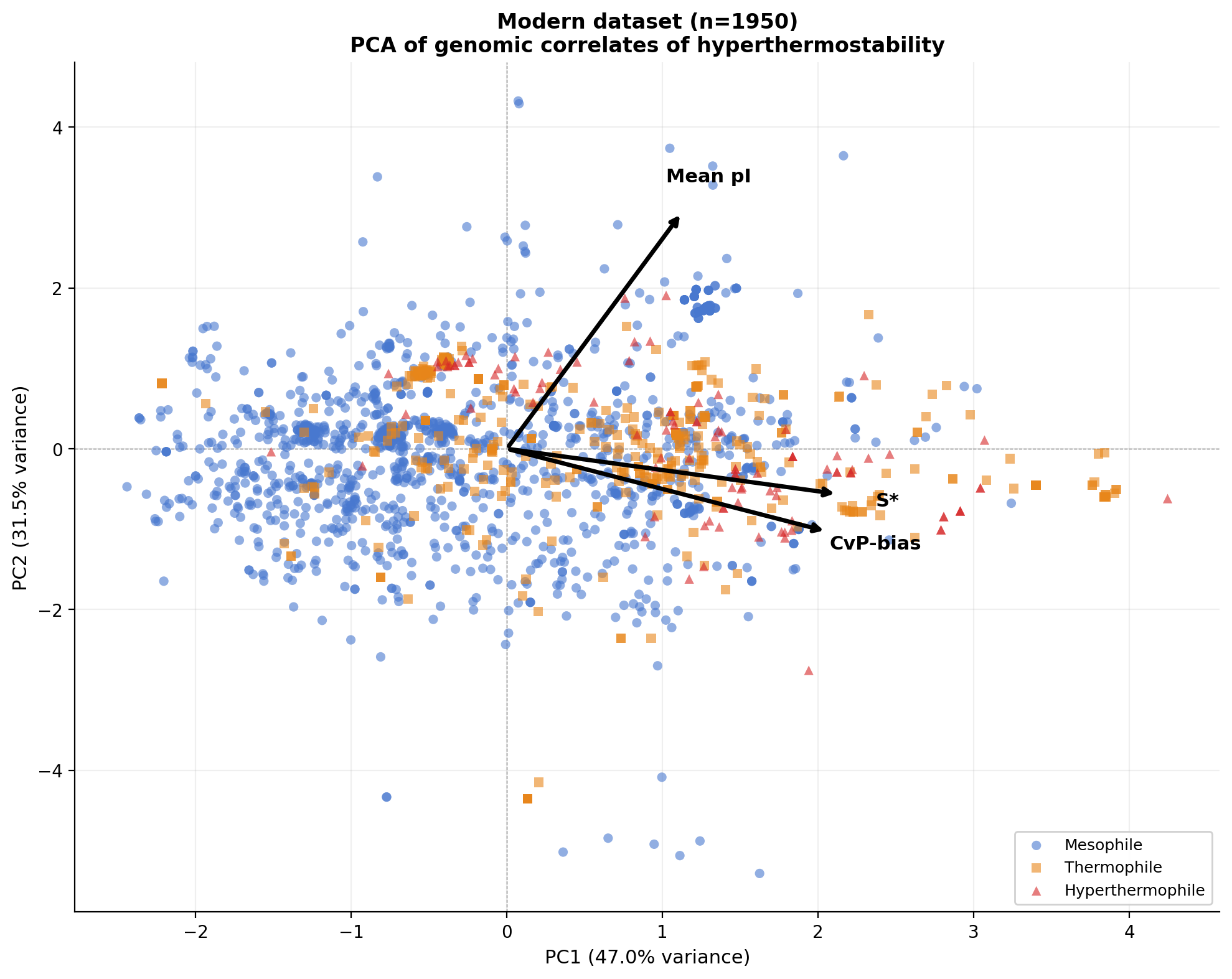

### figure2_original.png

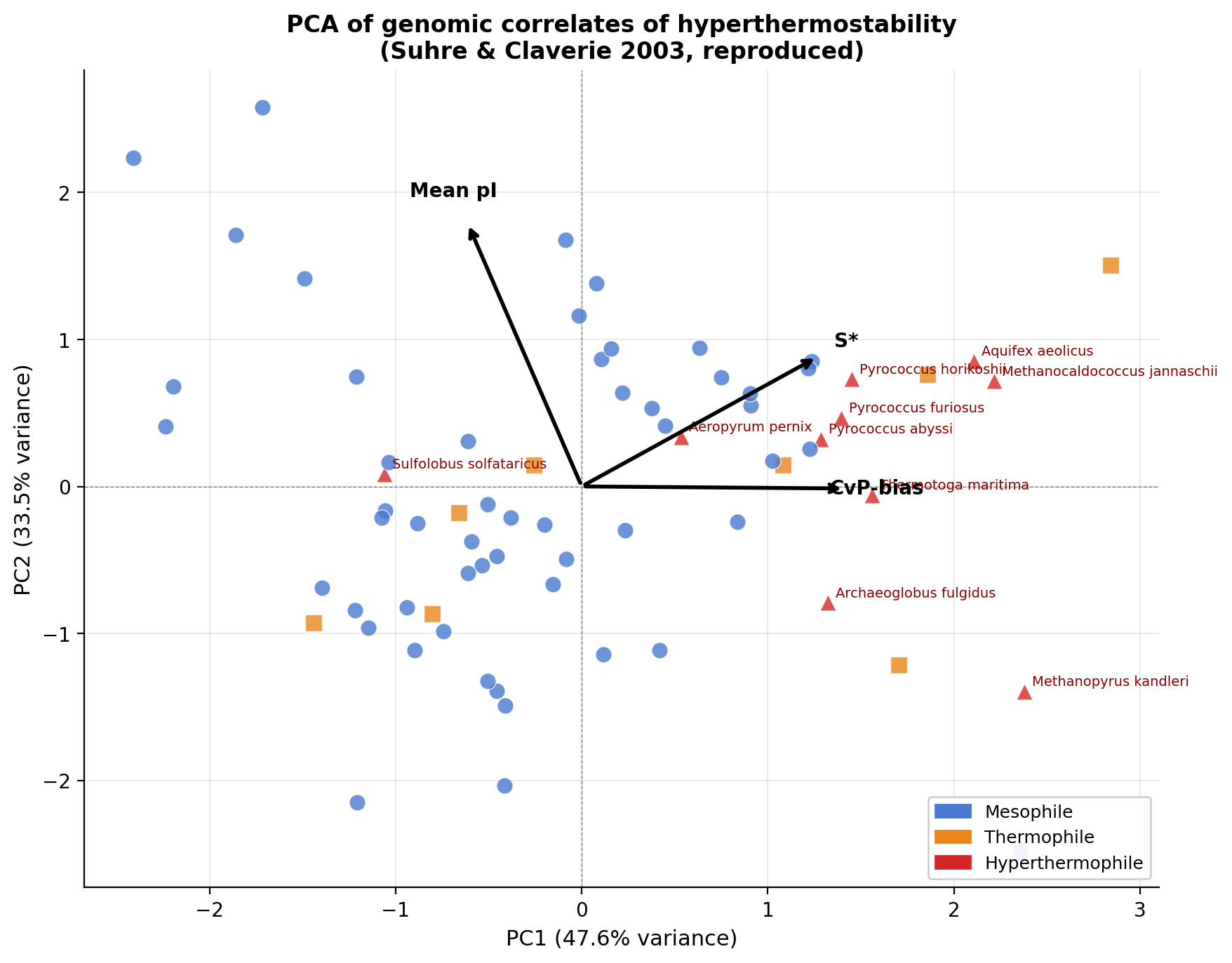

### figure3_cvp_vs_ogt_combined.png

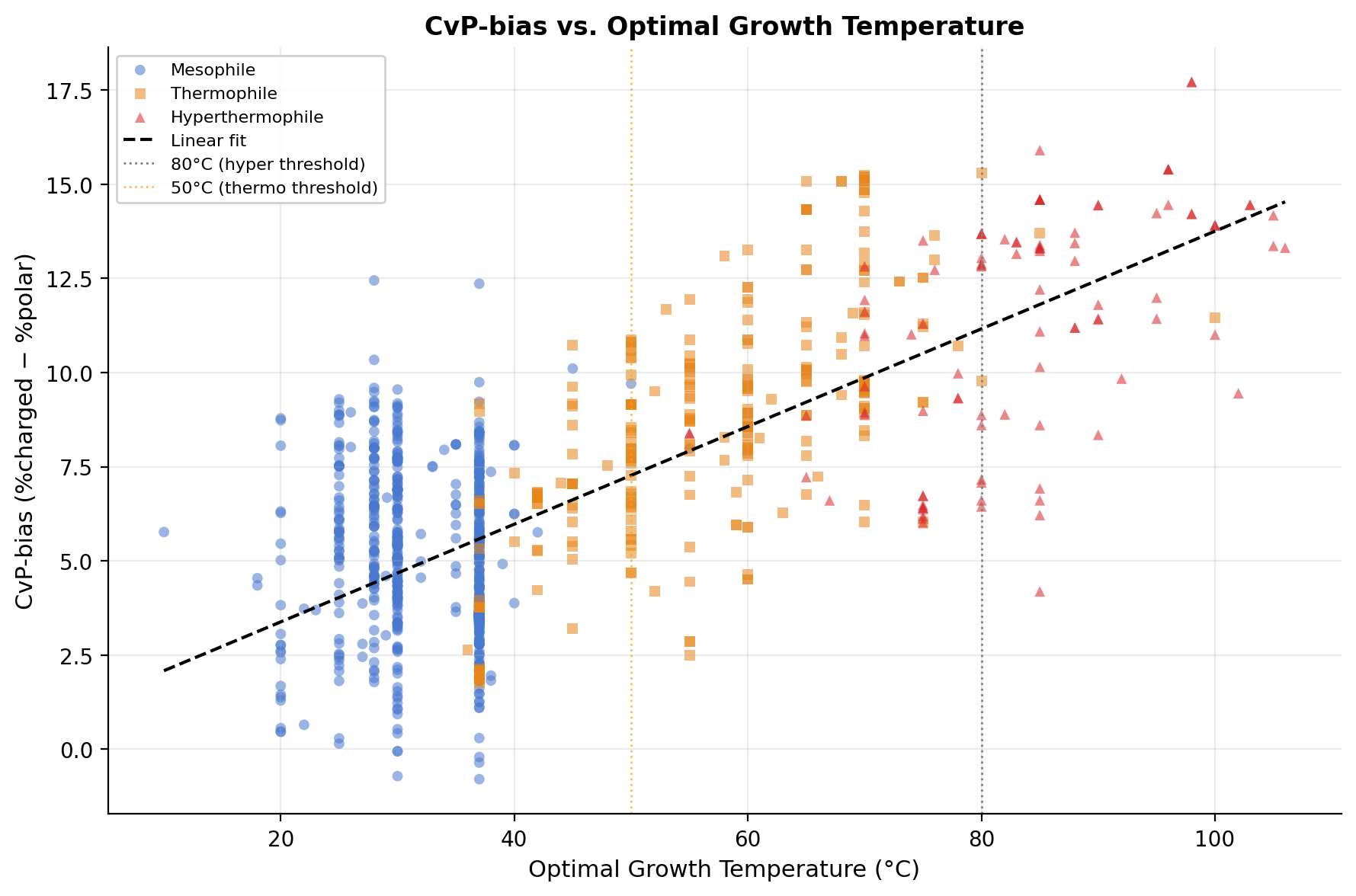
